# Epigenetic regulation of a heat-trainable small heat shock protein gene locus controls thermomemory in a unicellular alga

**DOI:** 10.64898/2026.03.31.715551

**Authors:** Elise Kerckhofs, Teresa Kowar, Martha R. Stark, Léa Faivre, Ruth Lintermann, Ana-Belen Kuhlmann, Stephen D. Rader, Daniel Schubert

## Abstract

A powerful way to enhance heat tolerance is to prime organisms with a moderate heat treatment to establish a molecular stress memory permitting the survival of the organism when exposed to subsequent heat shocks. While this has been extensively studied in multicellular organisms, we demonstrate that the unicellular red alga *Cyanidioschyzon merolae* exhibits heat stress memory. We show that, similarly to more complex organisms, thermomemory in this alga is underpinned by transcriptomic reprogramming, with the chloroplast emerging as the main site of gene trainability. Additionally, we find a conserved small heat shock protein (sHSP)-encoding locus in the nuclear genome to be heat-trainable, likely by histone depletion and sustained removal of the repressive mark histone H3 Lysine 27 trimethylation (H3K27me3). Of *C. merolae’s* two sHSPs, only the nuclear-localizing CmsHSP2 is necessary for proper HS memory establishment. Finally, we reveal a role for the H3K27me3-transferase CmE(z) (Enhancer of zeste) in heat stress memory which shapes the transcriptome to recurring heat exposures, beyond regulating the trainable sHSP locus. Overall, our work provides a molecular framework for the regulation of heat stress memory in a unicellular eukaryote.

## Introduction

Over the past decades, global warming has led to steadily increasing temperatures and more frequent heat stress (HS) events, posing a challenge for yields of agricultural crops and survival of wild plants and algae (Challinor et al., 2014, Zhao et al., 2017, Battisti and Naylor, 2009). At the cellular level, HS causes dehydration and protein denaturation and disrupts the integrity of membranes. Photosynthetic organelles are particularly susceptible to HS, as they can be a major site of excessive production of reactive oxygen species (ROS), harming photosynthetic efficiency and energy metabolism (Lípová et al., 2010, Tang et al., 2007, Nishiyama et al., 2011, Krumova et al., 2014).

While toxic effects of mild HS can often be counteracted by innate thermotolerance mechanisms (basal thermotolerance), prolonged exposure to severe HS is usually lethal. However, an organism’s threshold for heat lethality is not fixed, but thermotolerance can instead be trained and improved: if an exposure to a sublethal HS (priming HS) is followed by a stress-free lag phase, a molecular memory is formed that enables a more resilient response to future HS (Lämke and Bäurle, 2017, Balazadeh, 2022, Stipcich et al., 2024). This HS memory can lead to survival of otherwise lethal heat doses (Charng et al., 2006, de Klerk and Pumisutapon, 2008, Burke, 1998). Uncovering molecular mechanisms that underpin HS memory thus bears immense potential for agricultural exploitation.

Several effectors that enable enhanced thermotolerance have been identified in plants. These include heat shock proteins (HSPs) that can prevent protein misfolding and aggregation in a chaperone-like manner, as well as ROS-detoxifying proteins that reduce oxidative stress (Hartl et al., 2011, Vabulas et al., 2010, Rezayian et al., 2019). Accumulation of these proteins during recurring HS is a memory strategy that boosts thermotolerance and relies on heat-induced adaptations of the transcriptome (Larkindale and Vierling, 2008, Tomljanović et al., 2025, Oberkofler et al., 2021). Genes that respond differently to recurring heat events are termed memory genes. Their trainability can be achieved by modifications of the epigenome, as the level of chromatin compaction and the presence/absence of certain chromatin marks can influence the expression of associated genes (Mansisidor and Risca, 2022). FORGETTER1 (FGT1), for instance, interacts with the SwItch/Sucrose Non-Fermentable (SWI/SNF) complex to actively maintain low nucleosome occupancy and sustained accessibility to the transcription machinery for several HS memory genes in *Arabidopsis thaliana* (Brzezinka et al., 2016). Other chromatin-based regulations involve dynamic changes of histone mark patterns: stress exposure leads to histone 3 lysine 4 trimethylation (H3K4me3) of memory genes that can enable their sustained transcription for several days after the stress exposure has faded (Liu et al., 2018, Lämke and Bäurle, 2017, Friedrich et al., 2021, Pratx et al., 2023). This can build up a pool of HS effector transcripts, such as *HSP22.0* or *ASCORBATE PEROXIDASE2* (*APX2*) in Arabidopsis, which enables a faster reaction to a second heat exposure, allowing the plant to respond more resiliently. H3K4me3 is antagonized by the repressive mark H3K27me3. Upon heat exposure, H3K27me3 is removed from some HS trainable genes and not restored after return to non-stress conditions, which is thought to contribute to their sustained expression or faster reinduction during recurring HS (Liu et al., 2019a, Liu et al., 2014). However, a direct role for H3K27me3 has only been shown for a subset of H3K27-trimethylated genes, and further mechanisms that enable transcriptomic stress memory are not fully understood (Faivre et al., 2024, Yamaguchi et al., 2021).

Most studies of HS memory have focused on multicellular plants, and evidence of HS memory in unicellular phototrophs remains scarce. We therefore used the red alga *Cyanidioschyzon merolae* to investigate whether molecular HS memory is evolutionarily conserved in this divergent, unicellular phototroph. *C. merolae* belongs to the group of Cyanidiophyceae, one of the earliest-diverging red algal lineages, which emerged approximately 1.3 billion years ago (Yoon et al., 2006, Yoon et al., 2004). Their natural habitat in volcanic hot springs has driven exceptional adaptation to elevated temperatures, making Cyanidiophyceae well suited for studies of thermotolerance (Albertano et al., 2000, Ciniglia et al., 2004). *C. merolae* offers several experimental advantages over multicellular plants: it lacks a cell wall, divides rapidly, and possesses a compact haploid genome in which most genes are present in single copy (Nozaki et al., 2007, Matsuzaki et al., 2004, Fujiwara et al., 2021, Fujiwara et al., 2017, Kuroiwa et al., 1994, Ohnuma et al., 2008). These features facilitate genetic experiments while still retaining many conserved plant gene families and pathways, allowing insights into stress adaptation that may extend to higher plants (Misumi et al., 2005). Although chromatin-based stress regulation in *C. merolae* remains poorly characterized, the Polycomb repressive complex 2 (PRC2) is conserved, and H3K27me3 marks both transposable elements and gene bodies. This indicates that chromatin-mediated gene regulation and adaptive transcriptional plasticity are feasible in this alga (Mikulski et al., 2017, Hisanaga et al., 2023).

Here we demonstrate that *C. merolae* can be trained by heat priming to acquire enhanced thermotolerance. We show that this thermomemory is underpinned by transcriptomic reprogramming that is partially dependent on the H3K27 trimethyl transferase E(z) and is characterized by hyperinduction of chloroplast gene expression in previously primed cells. We further identify a heat-stress-trainable genomic locus encoding *C. merolae’s* two sHSPs whose trainability is mediated by histone eviction and concurrent loss of H3K27me3.

Interestingly, only the chloroplast-localizing sHSP is required for HS memory, while the nuclear-localizing one is dispensable. Overall, we reveal a molecular framework for the regulation of HS memory in *C. merolae* which is based on chromatin regulation of a trainable sHSP gene locus.

## Results

### Cyanidioschyzon merolae memorizes heat stress

To determine whether the unicellular red alga *C. merolae* can establish an enhanced HS tolerance after priming (thermomemory), we first defined lethal and sublethal HS conditions. *C. merolae* grows optimally at 42 °C, and a 60 °C HS has been reported to be lethal (Kobayashi et al., 2014). Consistent with this, incubation at 60 °C revealed survival for up to 40 min, whereas 1 h exposure was lethal, as assessed after five days of recovery at 42 °C (Fig 1A-B). We therefore defined 1 h at 60 °C as a lethal (triggering) HS.

**Figure 1:**
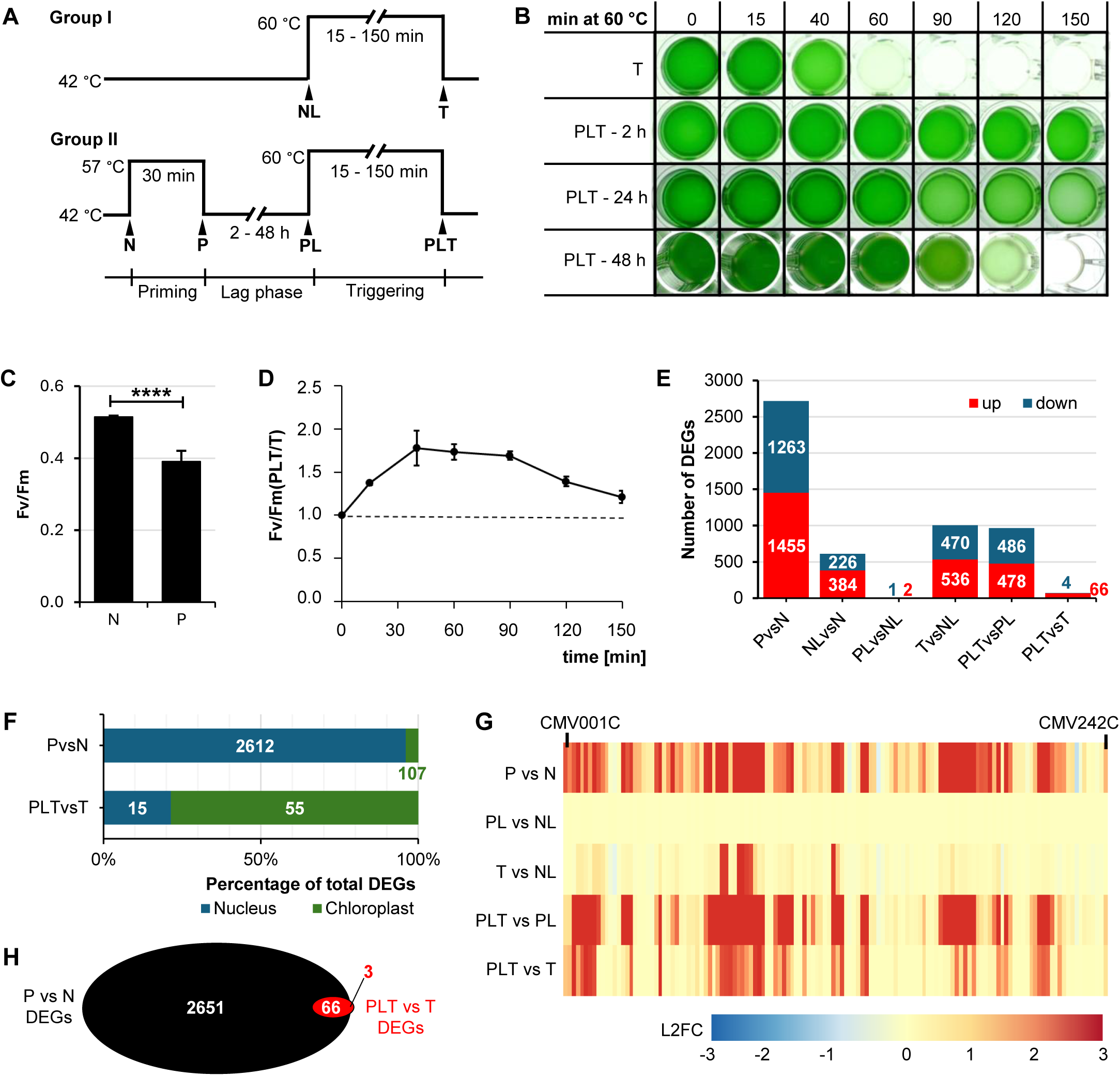
*Cyanidioschyzon merolae* establishes a powerful thermomemory upon HS priming. **A.** Experimental design. Algal cells were exposed to a triggering heat-stress (HS) at 60 °C either without prior priming (Group I, T), or after a 30 min priming HS at 57 °C, followed by a lag phase of 2, 24 or 48 h at 42 °C (Group II, PLT). **B.** Survival of unprimed (T) and primed (PLT) cells after 2 h, 24 h or 48 h lag phase, following up to 150 min at 60 °C and subsequent 5 d recovery at 42 °C. **C.** PSII quantum yields Fv/Fm of naïve (N) cells and cells primed for 30 min at 57 °C (P). Data show mean ± SD. Two-sided t-test, α = 0.05, n = 5 with three technical replicates per biological replicate, p < 0.0001 (****). **D.** Ratios of PSII quantum yields of primed vs unprimed cells Fv/Fm(PLT/T) after a 24 h lag phase and up to 150 min at 60 °C. Data show mean ± SD (n = 3 biological replicates with three technical replicates each). See Fig EV1 for untransformed Fv/Fm values. **E.** Total numbers of up-(red) and downregulated (blue) genes identified at different time points of the thermotolerance experiment (A). T and PLT samples were harvested after 15 min at 60 °C. Genes were considered differentially expressed if pl = -log(padj) x |L2FC| > 1.5 (n = 3 with three technical replicates per biological replicate). **F.** Genomic origin of DEGs identified in P vs N and in PLT vs T, indicated as percentage of total DEGs per comparison. Absolute DEG numbers are indicated on bars. A total of 4601 nuclear and 132 chloroplast genes were included in the analysis. **G.** Heatmap of log2 fold changes (L2FC) for chloroplast genes (*CMV001C* - *CMV242C*) across relevant comparisons. Colors indicate raw L2FC; 87 of 660 values were capped at +3 for better visualization.

We then tested whether a preceding sublethal priming (P) HS followed by a stress-free lag phase (L) could enhance tolerance to the triggering (T) HS. We identified 57 °C as a sublethal HS for *C. merolae*, with cells surviving up to 150 min (Fig S1). A priming treatment of 30 min at 57 °C followed by a 2 h lag phase at 42 °C markedly extended survival at 60 °C to 150 min (PLT, Fig 1A-B). Thus, HS priming powerfully boosts thermotolerance in *C. merolae*, demonstrating that HS memory is not restricted to multicellular organisms.

To assess the persistence of HS memory, we extended the lag phase to 24 or 48 h. A 24 h lag phase maintained survival for up to 150 min at 60 °C, whereas survival decreased to 120 min after a 48 h lag phase (Fig 1B). This indicates that HS memory in *C. merolae* starts to decline within 48 h and may be linked to the number of cell divisions.

Because HS rapidly impairs chloroplasts and photosystems (PS), leading to reduced quantum yields (Fv/Fm) (Lípová et al., 2010, Tang et al., 2007, Nishiyama et al., 2011, Krumova et al., 2014), we assessed photosystem performance by PAM fluorometry. Naïve (N) cells exhibited an Fv/Vm of 0.55 ± 0.02, indicating that 50% of the emitted light underwent photochemical energy conversion, while the remaining 50% was re-emitted (Fig 1C). Priming at 57 °C reduced Fv/Fm modestly to 0.4 ± 0.04. In contrast, exposure to the triggering HS (60 °C; T) caused a pronounced decline, with Fv/Fm values of 0.28 ± 0.01 and 0.18 ± 0.02 after 15 and 40 min at 60 °C, respectively (Fig 1C, EV1), indicating that the priming HS has only a limited impact on photosynthetic performance compared with the lethal stress.

To generate a relative measure of photosystem fitness, we formed Fv/Fm(PLT/T) ratios with Fv/Fm(PLT/T) = 1 indicating no priming benefit and Fv/Fm(PLT/T) > 1 indicating a corresponding increase in PS performance in primed algae. Strikingly, quantum yields of primed cells were almost two-fold higher than those of non-primed cells after 40 and 60 min at 60 °C, demonstrating that HS memory is beneficial to PS fitness in *C. merolae* (Fig 1D, EV1). Notably, the protective effect of priming on PS persisted for up to 150 min, which is consistent with the prolonged survival of PLT algae (Fig 1B).

### HS memory is orchestrated by extensive changes in gene expression levels

HS memory is frequently associated with global transcriptomic reprogramming that establishes a molecular memory of prior stress (Kappel et al., 2023, Olas et al., 2021). To test whether similar mechanisms operate in *C. merolae*, we performed comparative RNA-Seq on cells before (N) and after (P) priming, at the end of the lag phase (NL, PL) and after 15 min HS triggering at 60 °C (T, PLT, Fig 1A). To validate data reliability, we performed principal component analysis (PCA), hierarchical sample clustering and dispersion estimation. Biological replicates clustered together, confirming correct sample handling (Fig S2A-B). The first three principal components accounted for 54.2%, 15.1% and 11.8% of the variance, respectively, separating samples primarily by intensity of heat treatment (PC1), genotype (PC2), and experimental stage (PC3) (Fig S2A). Gene-wise dispersion estimates closely followed the fitted curve, supporting the reliability of the variance modelling (Fig S2C).

We then quantified differentially expressed genes (DEGs) during HS triggering. In unprimed cells (T vs NL), 536 genes were significantly upregulated and 470 were downregulated, while primed cells (PLT vs PL) showed 478 induced and 486 repressed genes, indicating comparable DEG numbers in both groups (Fig 1E). Direct comparison of primed and unprimed transcriptomes (PLT vs T) identified 70 HS memory genes, of which 66 were hyperinduced and only four were hyperrepressed in PLT vs T (Fig 1E, Table S1). Strikingly, genomic annotation revealed that only 15 of these genes were encoded in the nucleus, while the remaining 55 were chloroplast-encoded (Fig 1F, Table S2). Thus, more than 40% of the total 132 annotated chloroplast genes included in the analysis are HS trainable, identifying the chloroplast as a major site of transcriptomic thermomemory. Heatmap analysis of chloroplast gene expression showed that many genes were induced upon priming (P vs N), with expression levels returning to basal levels by the end of the lag phase (PL vs NL) and hyperinduction upon HS triggering in previously primed cells (PLT vs PL and PLT vs T, Fig 1G). This heat-trainable expression pattern likely contributes to enhanced thermotolerance by replenishing heat-damaged chloroplast proteins, consistent with the higher Fv/Fm values observed in primed cells during HS triggering (Fig 1D, EV1). Accordingly, genes encoding heat-sensitive PS core components such as D1 (*psbA*) and D2 (*psbB*) were among the hyperinduced memory genes (Table S2).

The chloroplast, and more specifically the photosynthetic electron transport chain, is a major site of increased ROS production during HS, due to excessive energy accumulation paired with decreased acceptor availability (Lípová et al., 2010, Tang et al., 2007, Nishiyama et al., 2011, Krumova et al., 2014). ROS can have harmful effects on proteins, thus hyperinducing genes that code for chloroplast proteins can be a strategy to counteract HS damage by supplying freshly synthesized protein.

Next, we analyzed DEGs in P vs N and in PL vs NL to determine how HS memory genes behave prior to secondary stress exposure. Consistent with observations in Arabidopsis and rice (Ling et al., 2018, Kushawaha et al., 2021), exposure to the priming HS elicited a stronger transcriptomic response than the triggering HS, with 1456 genes significantly upregulated and 1263 genes downregulated relative to naïve cells (Fig 1E)T. Thus, nearly half of all analyzed genes were heat-responsive, underscoring the extensive impact of HS priming on the *C. merolae* transcriptome. Genomic annotation revealed that a large fraction of chloroplast genes (107 out of 132 annotated) were heat-responsive (Fig 1F). The majority were upregulated in P vs N, further supporting chloroplast gene induction as a heat-protective mechanism. DEG overlay revealed that almost all (67) memory genes were also DEGs in P vs N (Fig 1H). We only identified three DEGs in previously primed algae by the end of the lag phase (PL vs NL): two induced (*CMP358C*, *CMC002C*) and one repressed (*CMT010C*). *CMT010C* was also repressed in PLT vs T, while neither of the other memory genes showed differential expression during the lag phase (Table S1). This indicates that memory gene hyperinduction is not caused by sustained induction throughout the lag phase, but rather by faster reinduction upon exposure to the secondary stress, suggesting a chromatin-based regulatory mechanism.

### A sHSP-encoding locus is HS trainable

Analysis of nuclear HS memory genes revealed that three genes (*CMJ099C*, *CMJ101C*, *CMJ102C*) are clustered on chromosome 10 (Table S1). Plotting normalized read counts across this locus showed that all genes *CMJ099C*, *CMJ101C*, *CMJ102C* and the previously non-annotated *CMJ100C* exhibit a similar expression pattern: induction during the priming HS, return to basal levels by the end of the lag phase, and hyperinduction in previously primed compared to unprimed cells upon secondary HS (PLT vs T; Fig 2A). Thus, all four genes constitute a HS-trainable locus. Notably, expression levels were higher in P than in PLT or T, likely reflecting the globally weaker transcriptomic response at 60 °C compared with 57 °C (Fig 1E), consistent with observations in plants (Ling et al., 2018, Kushawaha et al., 2021).

**Figure 2:**
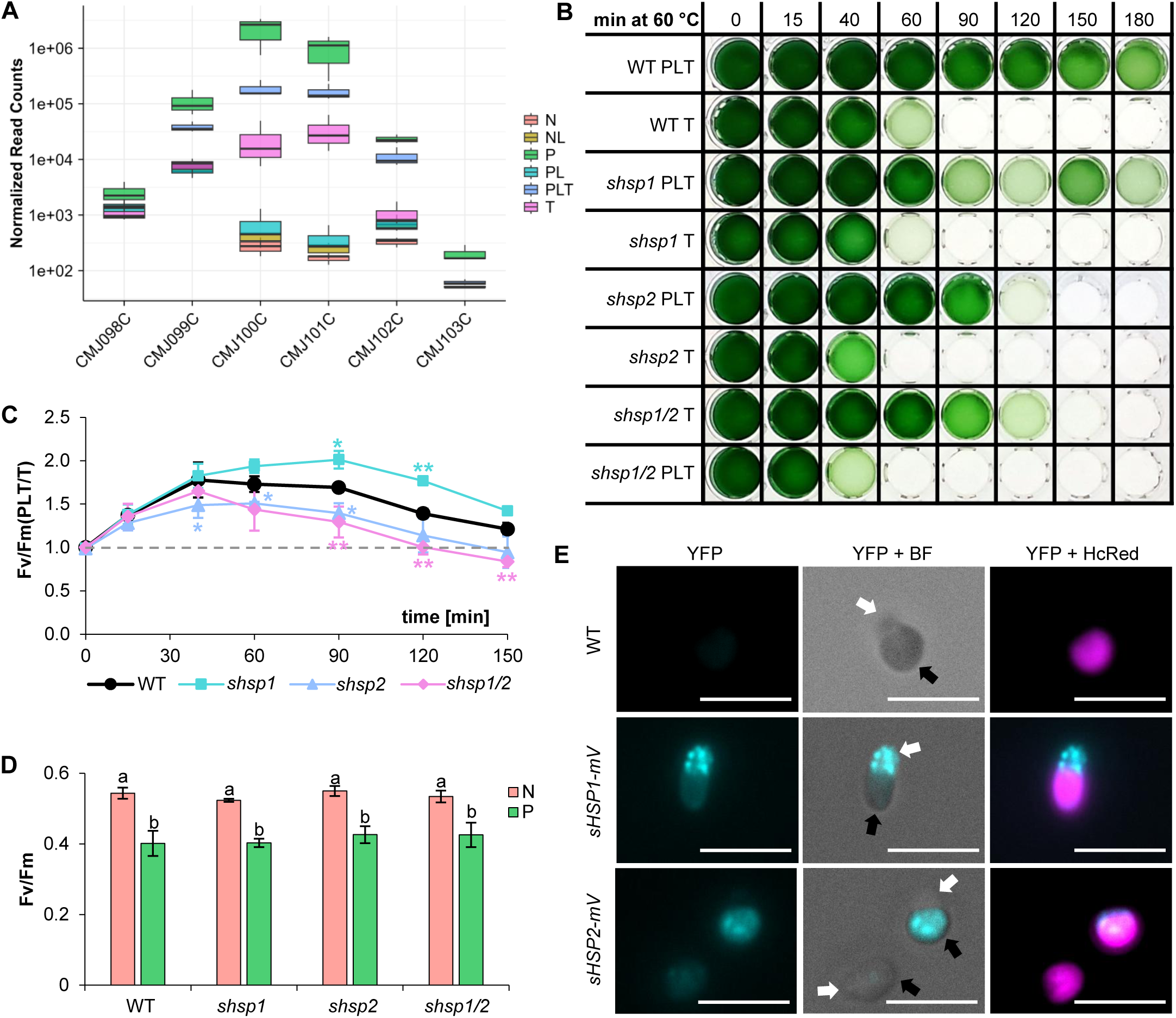
CmsHSP2 is a key effector of thermomemory in *C. merolae*. **A.** Log-transformed normalized RNA-seq read counts for genes within the memory locus *CMJ099C-CMJ102C* and flanking genes. T and PLT samples were harvested after 15 min at 60 °C. Data are shown as boxplots (n = 3). **B.** Survival of primed (PLT) and unprimed (T) sHSP deletion mutants (*CMJ100C* = *shsp1*, *CMJ101C* = *shsp2*) after up to 180 min HS triggering at 60 °C and subsequent recovery at 42 °C for five days (24 h lag phase). Representative plate from three independent experiments. **C.** Ratios of PSII quantum yields FvFm(PLT/T) of *shsp* mutants during HS triggering at 60 °C. Data show mean ± SD. Two-way ANOVA followed by Dunnet’s test for each time point, control group = WT, α = 0.05, n = 3 biological replicates with three technical replicates, * = p < 0.05, ** = p < 0.01, only significant p-values displayed. **D.** PSII quantum yields (Fv/Fm) before (N) and after (P) 30 min HS priming at 57 °C. Data show mean ± SD. Two-way-ANOVA followed by Tukey’s HSD test, n = 3 with three biological replicates per technical replicate, α = 0.05, different letters indicate significance with p < 0.05. **E.** Subcellular localization of sHSP1-mVenus sHSP2-mVenus. Cells were primed for 30 min at 57 °C, recovered for 1 h at 42 °C, and imaged by epifluorescence microscopy. Brightfield (BF), HcRed (chloroplast autofluorescence) and YFP (mVenus) channels are shown. Scale bar = 10 μm. Colors were changed to magenta (HcRed) and cyan (YFP) retrospectively for better visibility. Nuclear poles (white arrows) and chloroplast poles (black arrows) are indicated. Representative micrographs: see Fig EV3 for additional views.

Although flanking genes *CMJ098C* and *CMJ103C* were slightly heat-inducible, they were not HS trainable and define the boundaries of this memory locus (Fig 2A). *CMJ099C* encodes a formate-tetrahydrofolate ligase and *CMJ102C* encodes the mu subunit of adaptor-related protein complex 1 (Nozaki et al., 2007, Matsuzaki et al., 2004), neither of which has a known role in thermoprotection. In contrast, *CMJ100C* and *CMJ101C* encode *C. merolae’s* two sHSPs, CmsHSP1 and CmsHSP2, respectively. Given the established thermoprotective functions of sHSPs, their HS trainability suggested a role in HS memory. To test this, we generated single and double deletion mutants, *shsp1*, *shsp2* and *shsp1/2*, by homologous recombination and assessed thermomemory. Strikingly, while primed WT and *shsp1* cells survived 150 min at 60 °C, primed *shsp2* and *shsp1/2* cells survived only 90 min (Fig 2B), demonstrating that CmsHSP2 is necessary for HS memory, while CmsHSP1 plays a less important role. The identical phenotypes of *shsp1* and the double mutant further indicate that the two sHSPs do not function redundantly (Fig 2B).

We next assessed PSII thermotolerance and primability by measuring Fv/Fm. Compared with WT, both *shsp2* and *shsp1/2* exhibited reduced Fv/Fm ratios in primed versus unprimed cells starting at 40 min at 60 °C (Fig 2C, EV2). Nonetheless, PSII primability was not completely abolished, as Fv/Fm(PLT/T) remained above 1 for up to 90 min at 60 °C, indicating that additional mechanisms maintain high photosynthesis rates in previously primed cells (Fig EV2). Notably, *shsp1* displayed Fv/Fm(PLT/T) ratios comparable to WT for up to 60 min at 60 °C, and even higher values at later time points (Fig 2C, EV2), suggesting that CmsHSP1 is a negative effector of HS memory, which is somewhat contradictory to its likely heat-protective role and its hyperinduction in heat conditions.

Although both *CMJ100C* and *CMJ101C* were induced strongly during heat priming (Fig 2A), deletion of either or both had no effect on Fv/Fm after 30 min at 57 °C, indicating that these sHSPs are dispensable for basal photosystem thermotolerance (Fig 2C). Consistently, triggered-only (T) mutants did not differ from WT in Fv/Fm for up to 150 min at 60 °C (Fig EV3).

### *C. merolae’s* sHSPs localize to different organelles during HS

To further investigate the roles of the sHSPs in HS memory, we examined their subcellular localization during HS. Target peptide prediction tools suggested nuclear localization of CmsHSP1, while CmsHSP2 is predicted to localize to the chloroplast and/or mitochondrion (Kobayashi et al., 2014). To test this experimentally, we fused *CMJ100C* and *CMJ101C* to the yellow fluorescent protein (YFP) mVenus and stably integrated the construct into the *C. merolae* genome (Fig S3). Fluorescence microscopy of the resulting *sHSP1-mV* and *sHSP2-mV* strains revealed no detectable signal under non-stress conditions. Following induction by the 57 °C priming HS, mVenus fluorescence was detected in both strains 1 h after heat exposure (Fig 2E, EV3). In *sHSP1-mV* cells, the YFP signal was concentrated at the nuclear cell pole, indicating nuclear/perinuclear localization. In contrast, *sHSP2-mV* fluorescence co-localized with the chloroplast. These results confirm distinct subcellular localizations for the two sHSPs during HS, in agreement with previous predictions (Kobayashi et al., 2014), and further support non-redundant functions for CmHSP1 and CmHSP2, consistent with their differential contributions to HS memory (Fig 2B-D, EV2).

### The HS-trainable locus is regulated by histone depletion and H3K27me3

The shared expression patterns of *CMJ099C-CMJ102C* suggest a common regulatory mechanism, potentially involving chromatin modifications that control locus accessibility (Fig 2A). H3K27me3 and its antagonist mark H3K4me3 are known regulators of transcriptional HS memory in plants (Liu et al., 2018, Zeng et al., 2019, Faivre et al., 2024, Yamaguchi et al., 2021). Using previously generated H3K27me3 ChIP-Seq data from *C. merolae* (Mikulski et al., 2017), we identified *CMJ101C* as the only HS trainable gene carrying this repressive mark in naïve conditions, suggesting a potential regulatory role for this modification. To investigate this, we performed ChIP-PCR with antibodies binding to H3K27me3. We confirmed the presence of H3K27me3 at the promotor of *CMJ100C/CMJ101C* and across the complete coding region of *CMJ101C* in naïve cells (Fig 3A). Consistent with previous findings, we detected no H3K27me3 at the TSSs of *CMJ099C*, *CMJ102C,* and *CMJ103C,* demonstrating that only the center of the HS trainable locus is decorated with the repressive histone mark (Fig EV4).

**Figure 3:**
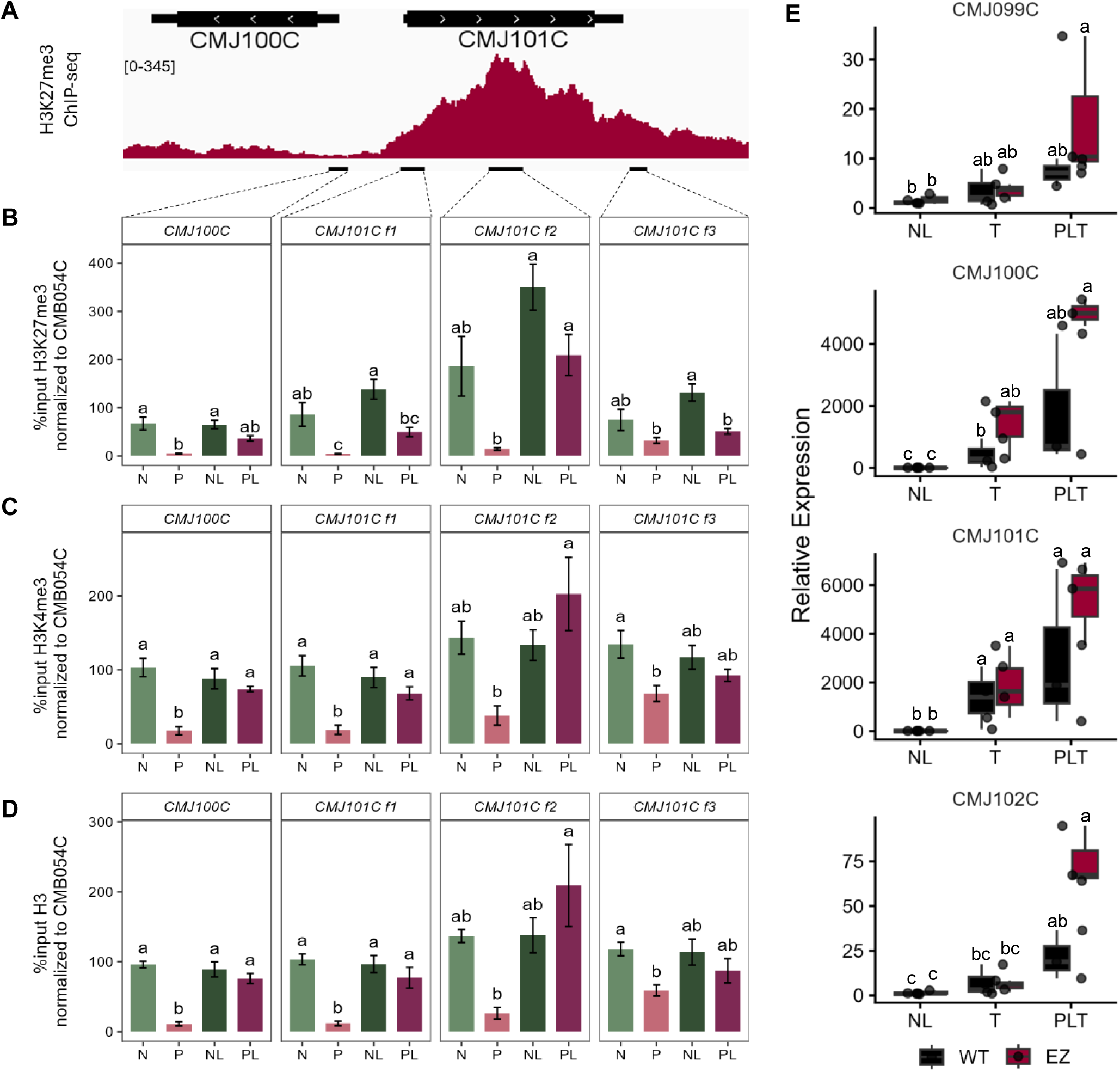
The sHSP memory locus is regulated by H3K27me3 and histone eviction. **A.** Genome browser view of H3K27me3 ChIP-Seq signal at the *CMJ100C-CMJ101C* locus, based on data from Mikulski et al. (2017). Black bars indicate regions analysed by ChIP-qPCR. **B-D.** Levels of H3K27me3 (**B**), H3K4me3 (**C**) and total H3 (**D**) across the *CMJ100C-CMJ101C* locus. Cells were grown at 42°C (N), primed for 30 min at 57 °C (P), or primed followed by a 24 h lag phase (PL). NL denotes unprimed controls harvested in parallel. ChiP-qPCR values are shown as % input normalized to the control locus *CMB054C*. Mean ± SEM, n = 4 biological replicates. One-way ANOVA followed by a Tukey’s post-hoc test (α = 0.05); different letters indicate significance with p < 0.05. **E.** Relative expression of *CMJ099C-CMJ102C* across memory conditions in WT and E(z) determined by RT-qPCR. Two-Way ANOVA followed by Tukey’s HSD test, α = 0.05, n = 3, different letters indicate significance with p < 0.05.

Following priming, H3K27me3 was almost completely lost from *CMJ100C* and *CMJ101C* (Fig 3B), consistent with their induction in P vs N (Fig 2A). This loss appeared specific to heat-trainable genes, as H3K27me3 levels at a heat-inducible but non-trainable gene (*CMG129C*) and at a heat-insensitive control gene (*CMB054C*) remained unchanged (Fig S5). Interestingly, total H3 occupancy was also significantly reduced after priming at *CMJ100C* and *CMJ101C* (Fig 3D). This suggests that histone depletion rather than active demethylation is the primary cause of H3K27me3 loss. Consistent with this, levels of the activating histone mark H3K4me3 were also reduced upon priming (Fig 3C), further supporting histone depletion as the mechanism for loss of methylation at this locus.

By the end of the lag phase, total H3 levels at *CMJ100C* and *CMJ101C* were restored and did not differ between primed (PL) and unprimed (NL) cells (Fig 3D, EV4D), indicating comparable nucleosome density. However, H3 levels were significantly lower in NL and PL cells compared to N and P for almost all genes tested (Fig 3D, EV4D, S5D). This might be caused by increased culture densities at the end of lag phase and is in line with the finding that 610 genes were differentially expressed in NL vs N, suggesting that the 24 h lag phase affects both the transcriptome and the chromatin (Fig 1E). Strikingly, the level of H3K27me3 was significantly lower in PL compared to NL at the TSS of *CMJ101C* (Fig 3B). A similar trend of lower H3K27me3 levels for previously primed cultures was observed downstream of its CDS and around the *CMJ100C* TSS. This indicates that re-incorporated nucleosomes are not fully re-methylated by the end of the lag phase, potentially facilitating faster reinduction upon secondary HS. In contrast, levels of the antagonizing H3K4me3 mark were consistent between NL, PL and N, suggesting that trainability of the memory locus is independent of H3K4me3 (Fig 3C, EV4C).

Together, these results support a model in which HS priming induces transcription-coupled histone depletion at the sHSP locus, leading to passive loss of H3K27me3. Partial persistence of this hypomethylated state after the lag phase may cause enhanced gene reinduction during triggering. We hypothesize that incomplete PRC2-mediated remethylation lowers the activation threshold and accelerates transcriptional reactivation during HS triggering. To test this, we analyzed relative expression levels of *CMJ099C-CMJ102C* in a previously described mutant that lacks the catalytic subunit of PRC2, *E(z)* (*CMQ156C*), and is completely devoid of H3K27me3 (Hisanaga et al., 2023). Strikingly, expression of all four genes was significantly higher in primed *E(z)* cells than in primed WT cells upon triggering (PLT), supporting the hypothesis that the absence of H3K27me3 at this locus by the end of the lag phase (PL) enables stronger induction of these genes upon exposure to the triggering HS (Fig 3E). Expression levels of *CMJ099C-CMJ102C* were similar between *E(z)* and WT in unstressed cells (NL), indicating that the absence of H3K27me3 enhances inducibility of the locus but is insufficient to activate transcription in the absence of heat stress. Furthermore, *CMJ099C-CMJ101C* expression was not enhanced in unprimed triggered (T) *E(z)* cells compared to WT (Fig 3E), suggesting that removal of H3K27me3 is not the only mechanism that controls heat trainability of this locus.

### E(z) is necessary for long-term HS memory

To test whether loss of *E(z)* affects thermotolerance, we subjected the mutant to the HS memory assay using a 2 h or 24 h lag phase (Fig 1A). Despite prior priming, *E(z)* cells only survived for up to 120 min at 60 °C after a 24 h lag phase, whereas WT cells survived 150 min, and PSII quantum yields, Fv/Fm(PLT/T), were significantly reduced in the mutant from 60 min at 60 °C onward (Fig 4A-B, EV5A). Thus, although sHSP transcripts are hyperinduced in the mutant (Fig 3E), HS memory is impaired rather than enhanced after a 24 h lag phase, indicating that PRC2 activity is required for full thermomemory.

**Figure 4:**
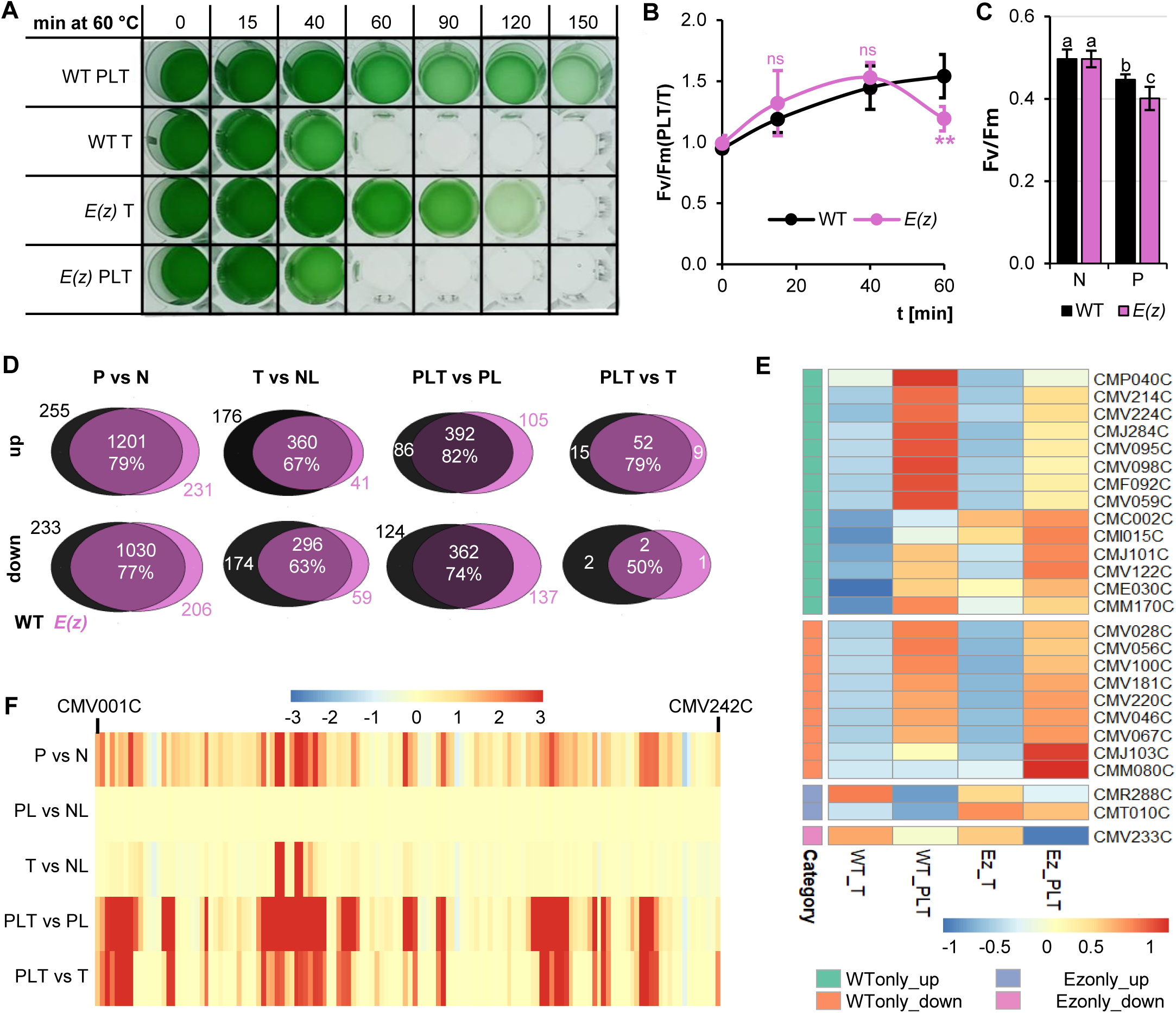
Loss of H3K27me3 compromises long-term heat stress memory. **A.** Survival of primed (PLT) and unprimed (T) cells after up to 150 min HS triggering at 60 °C and 5 d recovery at 42 °C (24 h lag phase). Representative plate of three independent experiments. **B.** Ratios of PSII quantum yields (Fv/Fm(PLT/T)) during HS triggering at 60 °C. Mean ± SD. Two-way ANOVA followed by Dunnet’s test for each time point (control group = WT, α = 0.05, n = 3 biological replicates with three technical replicates, ** = p < 0.01, ns = not significant). See Fig EV5A for untransformed Fv/Fm values. **C.** PSII quantum yields before (N) and after (P) 30 min HS priming at 57 °C. Mean ± SD. Two-way ANOVA followed by Tukey’s HSD test (α = 0.05, n = 3 biological replicates with three technical replicates, different letters indicate significance with p < 0.05. **D.** Overlay of DEGs identified in WT (black) and *E(z)* (pink) across thermomemory conditions. Percentages indicate shared DEGs. See Fig EV5B. **E.** Heatmap of z-scaled relative expression levels of PLT vs T memory genes unique to WT or *E(z)*. Chloroplast genes are indicated by “CMVxxxx” **F.** Heatmap of raw L2FC values for chloroplast genes (CMV001C - CMV242C) across thermomemory conditions. 60 of 660 values were capped at +3 for better visualization.

By contrast, we observed no differences between WT and *E(z)* when a short (2 h) lag phase was used, suggesting that PRC2 activity is dispensable for short-term memory but required for persistence over longer timescales, pointing to an epigenetic effect (Fig S5).

Notably, survival and PSII quantum yields of unprimed (T) *E(z)* cells exposed directly to 60 °C were indistinguishable from WT (Fig 4A, EV5A), suggesting that E(z)’s role in thermotolerance is specific to stress memory. However, we found that *E(z)* photosystems were more sensitive to the sublethal priming HS, as Fv/Fm declined more strongly in P vs N compared with WT (Fig 4C), indicating that PRC2 also contributes to basal thermotolerance at sublethal temperatures.

### HS memory-associated transcriptomic reprogramming partially depends on E(z)

The HS memory defect of *E(z)* cannot be explained by failure to induce the sHSP locus, as these genes were hyperinduced in the mutant (Fig 3E). Given that H3K27me3 marks ∼14% of the *C. merolae* genome under naïve conditions, including 242 coding genes and 172 repetitive elements (Mikulski et al., 2017), we reasoned that loss of E(z) likely affects HS memory through broader transcriptomic changes. We therefore performed RNA-Seq of heat-stressed *E(z)* cells and analyzed DEGs across the relevant conditions (Fig EV5B). During HS triggering, transcriptomic responses were largely preserved in the mutant: 61 genes were induced and three repressed in PLT vs T, similar to WT (Fig 4D). Of the 66 hyperinduced HS memory genes in WT, only 14 lost trainability in *E(z)* (Fig 4D, Table S2). This demonstrates that the transcriptomic response to the triggering HS is similar in both lines, and this limited loss of trainability likely explains why HS memory is not completely abolished in the mutant (Fig 4A-B).

Genes that lost trainability in *E(z)* included known or putative HS effectors, such as an L-asparaginase (*CMI015C*), a homologue of which was shown to be a HS chaperone in unicellular thermophiles (Dumina and Zhgun, 2023, Jena et al., 2018), and a cytochrome P450 (*CMJ284C*), whose homologue was shown to mediate HS tolerance in rice by balancing ROS homeostasis (Lv et al., 2025). However, closer inspection of read counts revealed that only four genes (*CMP040C*, *CMJ284C*, *CMF092C*, *CMM170C*) specifically failed to hyperinduce during PLT, whereas the other five genes (*CMC002C*, *CMI015C*, *CMJ101C*, *CME030C*, *CMM170C*) were already upregulated in triggered-only (T) *E(z)* cells and therefore classified as non-trainable (Fig 4E, S6). This indicates that H3K27me3 contributes to HS memory for a limited subset of effector genes, while regulating others independently of memory.

As in WT, the majority (56) of hyperinduced genes in *E(z)* were chloroplast-encoded (Table S2). Chloroplast gene expression dynamics were largely preserved, with induction during priming, partial relaxation during the lag phase, and hyperinduction during triggering (Fig 1G, 4F). This demonstrates that trainability of chloroplast genes is not affected in *E(z)* consistent with its role as nuclear chromatin regulator. Only seven chloroplast genes gained trainability in the mutant, further supporting that chloroplast trainability is largely independent of PRC2 (Fig 4E, Table S2). Nevertheless, *E(z)* cells showed reduced PSII fitness during HS memory and increased sensitivity to the priming HS (P vs N; Fig 4B). Together, this suggests that the photosynthetic efficiency of the mutant is more heat-susceptible than in WT (Fig 4B). This places the PRC2 complex as a potential upstream regulator of chloroplast heat resilience and might explain why the mutant’s ability to survive the 60 °C HS after priming was affected, considering that efficient photosynthesis is necessary for survival.

To further clarify the role of E(z) in HS memory, we compared transcriptomes across additional stages of the thermomemory assay. Following priming (P vs N), total numbers of up- and down-regulated genes were comparable between WT and *E(z)*, with substantial overlap (79% of induced and 77% of repressed DEGs; Fig 4D, EV5B). This suggests that loss of E(z) does not grossly alter the primary transcriptomic response to the priming HS.

Consistently, gene ontology (GO) enrichment analysis revealed similar functional categories among DEGs in both lines (Fig S7, Table S4). GO terms related to protein folding and translation were enriched among upregulated genes in both genotypes, reflecting activation of conserved heat shock response (HSR) pathways that counteract protein damage and enable proteome reprogramming (Alagar Boopathy et al., 2022, Swindell et al., 2007). The intact induction of these pathways in *E(z)* suggests that they are largely independent of H3K27me3 regulation in *C. merolae* and that PRC2 is not required for activation of the canonical HSR. Chloroplast- and photosynthesis-related GO terms were also enriched in both genotypes; however, enrichment strength and DEG ratios were consistently higher in WT, indicating that while chloroplast gene induction upon priming is retained in *E(z)*, it is less effective. This reduced induction is reflected in chloroplast gene expression heatmaps (Fig 1G, 4F) and likely contributes to the stronger decline in PSII quantum yields observed in *E(z)* during priming (Fig 4B).

Among repressed genes, top-enriched GO terms in both genotypes were related to membrane-associated processes and compartments, extracellular space and cell-cell signaling (Fig S7). This indicates that *C. merolae* represses energy-consuming secretory pathways and remodels lipid-based compartments that are naturally heat-sensitive due to their chemical properties. Overlap of enriched GO terms among repressed genes was lower between WT and *E(z)* than for induced genes, suggesting that PRC2 activity contributes more strongly to priming-associated transcriptional repression than to gene activation. This is in line with the repressive function of H3K27me3 and suggests that loss of PRC2 may subtly alter the balance between protective induction and adaptive repression during priming.

Comparison of transcriptional responses during the triggering HS further highlights this distinction. In primed cells (PLT vs PL), DEG overlap between WT and *E(z)* was high (82% induced, 74% repressed), whereas overlap was reduced in unprimed cells (T vs NL, 67% induced, 63% repressed) (Fig 4D). This suggests that H3K27me3 plays a larger role in shaping the basal HS response at 60 °C than in the transcriptional reprogramming associated with HS memory. Importantly, however, this transcriptional divergence in unprimed cells did not translate into altered survival or PSII performance (Fig 4A-B), demonstrating that PRC2-dependent transcriptional effects are not limiting for acute thermotolerance.

## Discussion

While molecular memory of environmental stress is well documented in multicellular organisms (Kishimoto et al., 2019, Roces et al., 2022, Hilker et al., 2016), evidence for such stress memory in unicellular organisms remains limited. Here, we show that the unicellular alga *C. merolae* can be primed by HS to acquire enhanced thermotolerance upon subsequent HS exposure. This demonstrates that molecular stress memory is not restricted to multicellular organisms and that it predates the appearance of metazoans, pointing to a highly conserved adaptive mechanism. HS memory was found to be sufficiently strong to permit survival of otherwise lethal heat doses, accompanied by a doubling of PSII quantum yields (Fig 1A-D). Beyond its evolutionary implications, this robustness highlights the potential translational relevance of HS memory mechanisms for crop improvement. Given its experimental tractability, *C. merolae* provides a powerful system to identify core regulators of stress memory that may be conserved across species.

As in higher plants, *C. merolae’s* thermomemory is underpinned by extensive transcriptomic reprogramming, reinforcing the idea that transcriptional plasticity is a conserved feature of HS memory and suggesting transcriptome modulators as key regulators (Liu et al., 2018, Liu et al., 2019b, Pratx et al., 2023, Lämke et al., 2016, Roces et al., 2022, Olas et al., 2021). Interestingly, we identified the chloroplast as the main site of transcriptomic thermomemory, with a large fraction of chloroplast-encoded genes being hyperinduced upon HS triggering in previously primed cells (Fig 1G). Several plant studies have demonstrated that the chloroplast is the most heat-sensitive organelle within phototrophic cells, as elevated temperatures rapidly disrupt photosystems and redox homeostasis, which can be harmful to proteins (Lípová et al., 2010, Tang et al., 2007, Nishiyama et al., 2011, Krumova et al., 2014). Hyperinducing chloroplast genes that code for members of the photosynthetic electron transport chain might thus be a strategy to exchange damaged proteins quickly, enabling sustained photosynthetic activity at unfavorable temperatures. While plants induce chloroplast and PS genes following a single HS exposure as part of the PSII repair cycle (Silva et al., 2003, Wang et al., 2018, Kato et al., 2018), our data extend this concept by identifying chloroplast gene hyperinduction as a hallmark of stress memory, rather than a transient stress response. This mechanism may thus represent an evolutionarily ancient strategy that could also contribute to HS memory in higher plants.

Among nuclear HS memory genes, we identified a heat-trainable locus comprising four adjacent genes (*CMJ099C-CMJ102C*), two of which encode the small heat shock proteins CmHSP1 and CmHSP2. sHSPs are highly conserved chaperones with well-established thermoprotective functions across all domains of life (Siddique et al., 2008, Waters and Vierling, 2020, Wi et al., 2023). Localization analyses revealed that CmHSP1 accumulates at the nuclear/perinuclear region, whereas CmsHSP2 localizes to the chloroplast during HS (Fig 2E), consistent with non-redundant roles in protecting distinct cellular compartments from stress damage. Functional analyses supported this division of labor: deletion of CmsHSP2 severely compromised PSII trainability and survival after recurrent HS, while deleting CmsHSP1 had no detrimental effect on photosynthesis rates (Fig 2C). These findings identify CmsHSP2 as a key effector of HS memory, likely acting within the heat-sensitive chloroplast to protect the proteome during HS in an ATP-independent, chaperone-like manner, analogous to plant chloroplast-localized sHSPs (Zhong et al., 2013, Bernfur et al., 2017).

The coordinated expression of *CMJ099C-CMJ102C* suggested shared regulation, and previous work showed that the TSS and gene body of *CMJ101C* are decorated with high levels of the repressive histone modification H3K27me3 under non-stress conditions (Mikulski et al., 2017), likely maintaining a compact chromatin state and gene silencing (Fig 5). Our chromatin analyses indicate that HS priming induces transcription-coupled histone depletion at this locus, leading to loss of H3K27me3 and transcriptional activation (Fig 3A-D). Importantly, while nucleosome occupancy is restored by the end of the lag phase, H3K27me3 is not fully re-established, leaving the locus in a more permissive chromatin state (Fig 5). This suggests that genes on this locus can be induced stronger in previously primed algae, because re-activation of transcription can occur faster due to the decreased load of repressive features (Fig 5). In line with this, we showed that deletion of the H3K27me3 writer E(z) led to even higher levels of *CMJ099C-CMJ102C* transcript in T and PLT conditions (Fig 3E).

**Figure 5:**
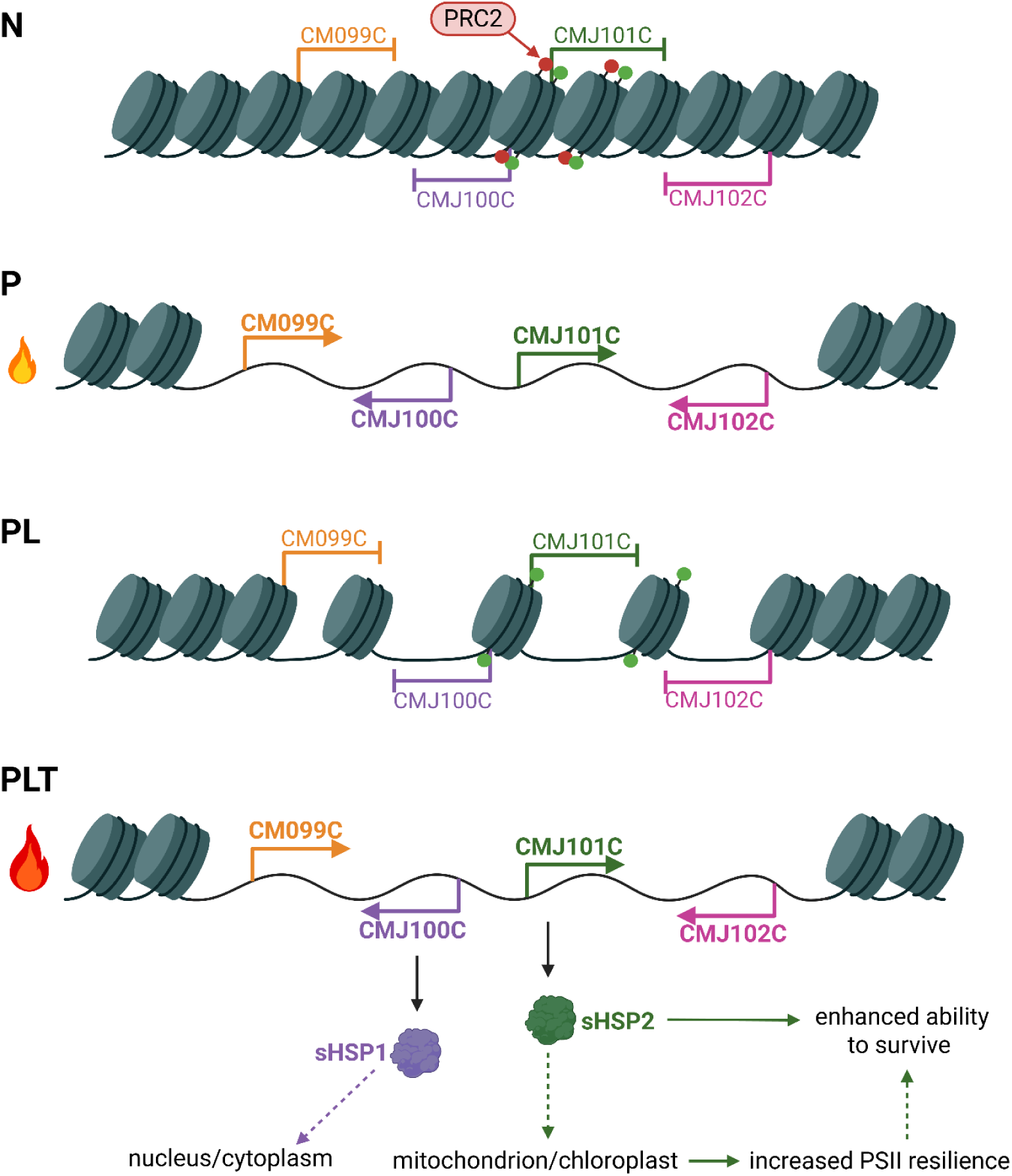
Model of HS-trainable sHSP locus regulation. In naïve (N) cells, the HS trainable locus *CMJ099C-CMJ102C* carries both activating K3K4me3 (green) and the repressive H3K27me3 (red) marks and adopts a condensed chromatin state that does not allow for gene expression. Deposition of H3K27me3 requires activity of E(z), the catalytic subunit of the PRC2 complex. HS priming (P) induces histone eviction, enabling transcription of *CMJ099C-CMJ102C*. By the end of the stress-free lag phase, histones and H3K4me3 marks are restored, but H3K27me3 remains absent, generating a permissive chromatin state that supports faster and stronger re-induction upon triggering HS. *CMJ100C* and *CMJ101C* encode CmsHSP1 and 2, respectively, which localize to distinct cellular compartments. CmsHSP2 promotes PSII resilience and survival, acting as a key effector of HS memory.

Notably, only *CMJ101C* encodes a protein with a demonstrated thermoprotective role, whereas the flanking genes appear dispensable for HS memory. We propose that their trainability reflects chromatin spillover from *CMJ101C*, as *CMJ100C* and *CMJ101C* are bidirectional genes that share a promoter, and the H3K27me3 enrichment was strongest at the center of the locus and gradually decreased towards its edges. Such regulatory spillover may be more common in compact genomes like that of *C. merolae,* which likely lack extensive insulator elements and chromatin boundaries found in higher eukaryotes.

Both H3K27me3 and its antagonist mark H3K4me3 are known to play a role in HS memory and gene trainability in plants, including sHSP loci (Yamaguchi et al., 2021, Liu et al., 2014, Pratx et al., 2023). However, we did not observe changes in H3K4me3 at this locus, suggesting that reliance on this modification for gene trainability may have evolved later in plant lineages. Future analyses of additional memory loci and histone methyltransferase mutants will be required to resolve this.

To test whether PRC2 activity contributes functionally to HS memory, we analyzed an *E(z)* mutant lacking H3K27me3. Despite hyperinduction of the thermotolerance-promoting *CMJ101C* in primed and triggered *E(z)* cells, we did not observe an improvement of thermomemory. Instead, the mutant was slightly compromised in memory establishment. Lack of a benefit of enhanced hyperinduction of *CMJ101C* for HS memory in *E(z)* could have several reasons: it is possible that CmsHSP2’s function in HS memory might not be dose-dependent, that transcript levels do not directly translate to protein levels, or that increasing CmsHSP2 levels might not be beneficial anymore after a certain threshold was exceeded (saturation). Interestingly, HS memory in *E(z)* was only affected after a 24 h lag phase, but not after a 2 h lag phase. This indicates that *C. merolae’s* PRC2 complex is needed for long-term rather than short-term memory, which is in line with PRC2’s reported role as an epigenetic mediator of long-term and trans-generational memory in plants (Gaydos et al., 2014).

Transcriptomic analyses revealed that loss of *E(z)* affected HS memory selectively. Only 15 genes lost trainability, many of which encode known or putative HS effectors. While most HS-responsive pathways remained intact, the mutant failed to properly regulate a few candidate memory effector genes, suggesting that PRC2 fine-tunes memory through targeted repression rather than global transcriptional control.

In summary, this study establishes *C. merolae* as a minimal and evolutionarily informative model for HS memory. We identify chloroplast gene hyperinduction and a trainable sHSP locus as central effectors of thermomemory and uncover a chromatin-based mechanism in which transcription-coupled histone depletion and delayed re-establishment of H3K27me3 enable enhanced gene reinduction. We show that PRC2 activity is required for long-term, but not short-term, HS memory, linking chromatin regulation by PRC2 to memory persistence in a unicellular system. Together, these findings argue that epigenetic stress memory is an ancient, conserved feature of eukaryotic life and provide a conceptual framework for dissecting core memory mechanisms that may be exploitable to enhance stress resilience in crops.

## Methods

### Algae material and growth conditions

**A.** *C. merolae* 10D strain (NIES-3377) was obtained from the Microbial Culture Collection at the National Institute for Environmental Studies in Tsukuba, Japan and maintained in 2x Modified Allen’s (MA2) medium at room temperature and continuous shaking at 50 rpm (Minoda et al., 2004). For experimental preparation, algae were grown at 42 °C and constant light (90 µE) and ambient air supply for several days.

### Generation of knock-out mutants

Generation of the H3K27me3-deficient mutant *E(z)* by homologous recombination using the *URA5.3* selection marker was previously described (Hisanaga et al., 2023). The CmsHSP1/2 single and double deletion mutants *shsp1*, *shsp2* and *shsp1/2* were generated by homologous recombination using the *chloramphenicol acetyltransferase* (*CAT*) resistance marker. Approximately 500 bp 5’ and 3’ homology arms flanking the *CmsHSP1* (*CMJ100C*) and *CmsHSP2* (*CMJ101C*) coding regions were amplified from genomic DNA using the high-fidelity DNA Polymerase SuperFi II (Thermo Fisher Scientific) and primer pairs 1 + 2 (*5’-CMJ100C*), 3 + 4 (*3’-CMJ100C*), 5 + 6 (*5’-CMJ101C*) and 7 + 8 (*3’-CMJ101C*), each carrying 15 bp overlaps for ligation independent cloning (LIC) into SwaI (5’ arm) and PacI (3’ arm) sites (Table S5; Scheich et al., 2007). PCR products were gel-purified on a 1% Agarose gel in 1 x TAE buffer (40 mM Tris base, 20 mM acetic acid, 1 mM EDTA), stained with 0.2 µg/ml ethidium bromide, and extracted using the Monarch Spin DNA Gel Extraction kit (New England Biolabs). Purifed homology arms were treated with T4 DNA Polymerase (Thermo Fisher Scientific), in the presence of 1.25 mM dCTP (5’ homology arm) or 1.25 mM dGTP (3’ homology arm) to generate complementary 15 bp overhangs for LIC. The backbone plasmid pSR886, carrying the *CAT* resistance cassette, LIC cloning sites, and an Ampicillin (Amp) resistance gene for transformation of *E. coli* was digested with SwaI (Thermo Fisher Scientific) and subsequently treated with T4 DNA polymerase in the presence of 1.25 mM dGTP. Linearized vector (0.01 pmol) was incubated with the 5’ homology arm (0.02 pmol) for 10 min at room temperature and transformed into chemically competent *E. coli DH10B* cells (Inoue et al., 1990). Following plasmid isolation and restriction enzyme digestion validation, constructs were digested with PacI, treated with T4 DNA ligase in the presence of 1.25 mM dCTP, and incubated with the corresponding 3’ homology arm to generate plasmids pEK01-1 for deletion of *CMJ100C* (*shsp1*) and pEK01-2 for deletion of *CMJ101C* (*shsp2*). To generate the double mutant *shsp1/2*, the 5’ homology arm of *CMJ100C* and the 3’ homology arm of *CMJ101C* were cloned sequentially into pSR886 using the same LIC strategy, yielding plasmid pEK02. Linear DNA fragments for stable genome integration were generated by PCR using primer pairs 9 + 10 (pEK01-1, *shsp1*), 11 + 12 (pEK01-2, *shsp2*), and 9 + 12 (pEK02, *shsp1/2*; Table S5).

Polyethylene glycol (PEG)-mediated transformation of *C. merolae* 10D was performed as described previously (Fujiwara et al., 2017, Villegas-Valencia et al., 2025) with minor modifications. Cells were grown in MA2 supplemented with 50 mM Glycerol (MA2G) to an OD_750_ of 1.2, washed with pre-warmed (42 °C) MA-I. Cell pellets were resuspended in an equal volume of MA-I. For each transformation, 25 µl of cells were mixed with 1 pmol of linear DNA in MA-I and 60 µg boiled, sonicated salmon sperm DNA. Subsequently, 125 µl of 60% PEG-4000 in MA-I was added, and the mixture was rapidly mixed and diluted into 40 ml MA2. Cells were incubated for 24 h at 42 °C under constant light (90 µE) and 5% CO_2_. Transformants were selected in MA2 supplemented with 200 µg/ml chloramphenicol for up to two weeks. Fresh chloramphenicol was added every two to three days and the medium was exchanged after seven days. After two weeks, random dilutions of cells were plated on solidified MA2G plant agar containing 200 µg/ml chloramphenicol to isolate single colonies.

### Generation of mVenus-tagged sHSP mutants

To track the subcellular localization of CmsHSP1/2, *CMJ100C* and *CMJ101C* were fused in frame to the yellow fluorescent protein mVenus and stably integrated into the *C. merolae* genome by homologous recombination. As 5’ homology arms, the coding sequences of both genes were amplified from genomic DNA using primer pairs 13 + 14 (*CMJ100C*) and 15 + 16 (*CMJ101C*), respectively (Table S5). Primers introduced BamHI and SwaI restriction sites and were designed to exclude the stop codon, enabling in-frame fusion with mVenus. The amplicons were gel-purified as described above and blunt-end cloned into the pJet intermediate vector using the cloneJet kit (Thermo Fisher Scientific). Constructs were transformed into chemically competent *E. coli* as described (Inoue et al., 1990), and after plasmid purification and validation, inserts were excised using BamHI and SwaI and gel-purified. The backbone vector, plasmid pSR1026, carrying *mVenus*, a flexible GSGS linker, a *CAT* resistance cassette, and multiple cloning sites, was digested with BamHI and SwaI and gel-purified. The purified 5’ homology arms were ligated into the linearized pSR1026 using T4 DNA ligase (Thermo Fisher Scientific) according to the manufacturer’s instructions and transformed into *E. coli DH10B*. Following validation, plasmids were digested with PacI, gel-purified, and treated with T4 DNA Polymerase in the presence of 1.25 mM dCTP to generate 15 bp overhangs for LIC. To generate 3’ homology arms, ∼ 500 bp downstream of the respective coding sequences were amplified from genomic DNA using primer pairs 17 + 18 (*CMJ100C*) and 19 +,20 (*CMJ101C*) each introducing complementary 15 bp LIC overhangs. Amplicons were gel-purified and treated with T4 DNA Polymerase in the presence of 1.25 mM dGTP. LIC was performed as described above, yielding plasmids pEK21 for generation of *sHSP1-mV* and pEK22 for generation of *sHSP2-mV*.

Linear DNA fragments for stable genome integration were generated by PCR using primers 21 + 22 (pEK21, *sHSP1-mV*), and 23 + 24 (pEK22, *sHSP2-mV*; Table S5). Transformation of *C. merolae* 10D with subsequent chloramphenicol selection was performed as described above.

### Heat stress experiments, photosystem fitness, and growth assays

Cells were grown at 42 °C to an OD_750_ of 1.0. For heat priming (see Fig 1A), cells were incubated in a 57 °C water bath with closed lid for 30 min (P). A control group was kept at 42 °C (N). After priming, both cultures were grown at 42 °C, constant light (90 µE) and ambient air supply for 2, 24 or 48 hrs. Both previously primed (PL) and unprimed cells (NL) were subjected to the triggering HS of up to 150 min at 60 °C in a water bath with closed lid. Primed and unprimed cells (T) were sampled at different time points (see Fig 1A for setup). To assess photosystem fitness of heat-stressed cells, PSII quantum yields were determined by Pulse Amplitude Modulation (PAM) Fluorometry using a PAM IMAG-K4B Fluorometer (Heinz Walz GmbH). Cells were dark-adapted for 8 min in 50 µl triplicate in 96-well plates, and Fv/Fm were determined at the following settings: Intensity 3, Frequency 1, Gain 1, Damp 2, Saturation Pulse 5. To assess their ability to survive different HS exposures, stressed cells were diluted to an OD_750_ of 0.1 in 48 well plates in MA2 (1 ml per well). Plates were incubated for five days inside plastic containers at 42 °C with 5% CO_2_ and constant light (90 µE) supply and then photographed to determine survival rates.

### RNA extraction and RT-qPCR

Cells were heat-stressed as described above. For RNA preparation, 2 ml of cells were centrifuged at 3,500 x g at 4 °C for 3 min. RNA was extracted using the innuPREP Plant RNA Kit (Analytikjena), adding 1% β-Mercaptoethanol to the lysis solution to increase RNA quality. DNase treatment and cDNA synthesis were performed using DNase I, RNase-free (Thermo Scientifc) and the RevertAid First Strand cDNA synthesis kit (Thermo Fisher Scientific) with oligo(dT) primers: 43 + 44 (*CMJ099C*), 45 + 46 (*CMJ100C*), 47 + 48 (*CMJ101C*), 49 + 50 (*CMJ102C*), 51 + 52 (*CMS262C*), 53 + 54 (*CMN304C*), 55 + 56 (*CMM193C*,Table S5). RT-qPCR was performed with 10 ng cDNA using the TakyonTM ROX SYBR 2X MasterMix dTTP blue (Eurogentec) according to the manufacturer’s instructions on the QuantStudio5 (Applied Biosystems). Temperature cycle program: 1x 3 min at 95 °C; 40 x 3 s at 95 °C followed by 30 s 60 °C; 15 s at 95 °C; 1 min at 60 °C, 15 s at 60 °C. Relative expression levels were determined according to the ΔΔCt method (Pfaffl, 2001) with mean Ct values of three reference genes CMS262C, CMN304C, and CMM193C. For each primer pair binding efficiencies were calculated based on a standard curve generated from 1:1, 1:10, and 1:100 dilutions of a cDNA pool. Efficiencies between 90 and 110 % in combination with a regression factor 0.95 < R^2^ were considered suitable.

### RNA-Seq

For RNA-Seq, RNA was extracted from heat-stressed cells in biological triplicate and treated with DNase I, RNase-free (Thermo Fisher Scientifc) as described. Library preparation and sequencing were conducted by Novogene UK (Cambridge). Libraries were prepared using polyA enrichment and the NEB Ultra RNA Library Prep Kit to produce unstranded data. Sequencing was performed on an Illumina NovaSeq 6000 platform, resulting in 150 bp long paired-end reads and 3G of raw data per sample.

### Bioinformatic analysis of sequencing data

Bioinformatic analyses were performed using Curta, the High Performance Comput of Freie Universitaet Berlin (Bennett, 2020). Raw sequencing read quality was validated by FastQC v0.11.9 (Andrews, 2010). Reads were aligned to the *C. merolae* genome using the ENSEMBL genome sequence and genome annotation (Cunningham et al., 2022) and STAR aligner v2.7.9a (Dobin et al., 2013). More than 92.5% of the reads were mapped for each sample. SAM files were converted into BAM files and indexed with SAMtools v1.11 (Li et al., 2009). Reads were counted with FeatureCounts v2.0.1 (Liao et al., 2014) after exclusion of low-quality reads (MAPQ < 10). For visualization of read coverage, BAM files were converted to BigWig format using DeepTools v3.3.2 (Ramírez et al., 2016) and inspected in the Integrative Genomics Viewer (IGV) v2.10.3 (Robinson et al., 2011). Differential gene expression analysis was performed in R v4.5.1 using DESeq2 v1.48.1 (Love et al., 2014), and a design formula that includes genotype, condition and their interaction (∼genotype x condition). Genes with fewer than 100 counts in at least two biological replicates were excluded. Sample clustering was investigated by principal component analysis (PCA) and sample distance mapping. Size-factor normalization and dispersion estimation for relevant comparisons were performed using DESeq2 default procedures, and model fitting was conducted using a local dispersion fit. To improve effect size estimation for low count genes, L2FC were shrunk using the ashr method (Stephens et al., 2023). Adjusted *p*-values (padj) were calculated using the Benjamini–Hochberg (Benjamini and Hochberg, 1995). To determine significant DEGs, pl ratios (pl = -log(padj) x |L2FC| > 1.5) were calculated. DEGs were considered significant if pl > 1.5. Normalized read counts across conditions and overlap of DEG sets (Venn diagrams) were generated using the ggplot2 v4.0.1 package (Wickham, 2016). For heatmaps, raw L2FC (chloroplast genome) or per-gene-z-scaled normalized counts (DEGs) were visualized using the pheatmap package (Kolde, 2025).

Gene Ontology (GO) enrichment analysis was performed with the topGO package v2.60.1 (Alexa and Rahnenfuhrer, 2025) and an annotation file that was generated from the QuickGO and Uniprot *C. merolae* GO annotations (The UniProt, 2025, Binns et al., 2009). The analysis was limited to protein-coding genes that were annotated with at least one GO term to avoid annotation bias. Proportions of included genes per gene set are stated in figure titles. Analyses were conducted separately for Biological Process (BP), Cellular Component (CC) and Molecular Function (MF) ontologies. Enrichment significance was assessed using Fisher’s exact test, applying the classic, elim, and weight01 algorithms, with the topology-aware weight01 method used for ranking and interpretation of enriched terms. GO terms with fewer than five annotated genes were excluded. p-values were adjusted using the Benjamini-Hochberg method (padj).

### Chromatin Immunoprecipitation (ChIP)

The chromatin immunoprecipitation (ChIP) protocol was adapted from Mikulski et al. (2017). Briefly, 10 mL of culture were crosslinked in 1% (v/v) formaldehyde for 10 min, after which glycine was added to a final concentration of 125 mM followed by a 5 min incubation. TritonX-100 was added to a concentration of 0.01% (v/v) to facilitate pelleting of the algae and the cells were washed twice with ice cold PBS buffer supplemented with 0.01% TritonX-100. In between washes, cells were pelleted by centrifugation at 5000g, 4°C for 10 min. After the washes, the cells were resuspended in 500 µL of extraction buffer (50 mM Tris-HCl pH 8, 10 mM EDTA, 1% SDS). The samples were then sonicated for 5 cycles (30 s ON, 90 s OFF) using a Bioruptor device (Diagenode). Subsequent steps were performed as described in Faivre et al. (2024), using α-H3K27me3 (C15410195 Diagenode), α-H3K4me3 (C15410003, Diagenode) or an α-H3pan (C15200011 Diagenode) antibodies. The qPCR was performed using the Takyon ROX SYBR MasterMix blue dTTP kit and QuantStudio5 (Applied Biosystems). The primers used for analysis are listed in Table S5 (25 - 42). Data is expressed as percent input normalized to the control locus *CMB054C* (% input GOI / % input CMB054C * 100). *CMB054C* was selected as the control locus due to its constant expression in all tested conditions and presence of H3K27me3.

### Fluorescence Microscopy

To investigate the subcellular localization of the fluorescently tagged proteins, cells were kept at 42 °C (control) or heat-stressed at 57 °C for 30 min and subsequently recovered at 42 °C for 1 h to allow protein maturation. All fluorescence microscopy was carried out with the SBS Canterbury Zeiss AxioImager.M1 fluorescence microscope in brightfield (BF) and fluorescence channels using HcRed (excitation wavelength 580 – 604 nm, emission wavelength 625 – 725 nm, exposure time 3.1 ms) and YFP (excitation wavelength 450 – 490 nm, emission wavelength 500 – 550 nm, exposure time 600 ms) filter sets, with illumination intensity set to 30%, and at 100 x magnification (acquisition software: ZEISS ZEN blue edition). Cells were visualized without prior staining or fixing to maximize mVenus signal quality. Representative images were exported as .tiff files and processed in ImageJ v1.54g (Schneider et al., 2012). Red channel (HcRed) was converted to magenta and green channel (YFP) was converted to cyan manually to increase visibility. Brightness and contrast adjustments were made according to the settings listed in Table S3.

### Vector sequences

See Supplementary File S1.

## Data availability

The datasets produced in this study are available in the following databases: RNA-Seq data: [paste link here]

## Acknowledgements

SDR acknowledges support from an NSERC Discovery Grant and from UNBC’s Office of Research and Innovation. DS, EK and TK acknowledge support through DFG grant CRC973. TK was funded by an Elsa-Neumann-Fellowship. Open Access Funding is provided by Freie Universität Berlin.

## Expanded View Figure Legends

**Figure EV1: PSII quantum yields during HS triggering.**

Cells were treated according to Fig 1A (24 h lag phase) c, red) and unprimed (T, black) cells were determined by PAM fluorometry. Plot shows Fv/Fm prior to Fv/Fm(PLT/T) transformation in Fig 1D. Mean ± SD. One-way ANOVA followed by Tukey’s HSD test, n = 3 biological replicates with three technical replicates, α = 0.05, p < 0.05 (*).

**Figure EV2: Fv/Fms of primed and unprimed *shsp* deletion mutants.**

Untransformed PSII quantum yields Fv/Fm of primed (PLT) and unprimed (T) WT and *shsp*. Cultures were treated as described (Fig 1A) with a 24 h lag phase. Mean ± SD (n = 3 biological replicates with three technical replicates). For transformed data Fv/Fm(PLT/T) see Fig 2C.

**Figure EV3: Subcellular localization of *sHSP1-mV* and *sHSP2-mV*.**

Micrographs of *sHSP1-mV* and *sHSP2-mV*. Cells were kept at 42 °C or heat-stressed for 30 min at 57 °C, and recovered at 42 °C for 1 h, before analysis with an epifluorescence microscope in brightfield (BF), HcRed (chloroplast autofluorescence) and YFP (mV) channels. Colors were changed to magenta (HcRed) and cyan (YFP) retrospectively for better visibility. Scale bar = 10 μm. Nuclear poles (white arrows) and chloroplast poles (black arrows) are indicated. Images of mutants after HS are represented in duplicates (1, 2).

**Figure EV4: Histone marks of *CMJ099C*, *CMJ102C* and *CMJ103C*.**

**A.** Genome browser view of H3K27me3 ChIP-Seq signal based on the data from Mikulski et al. (2017). Black bars indicate the location of the fragments examined in the ChIP-qPCR data presented below. **B. - D.** Levels of H3K27me3 (B), H3K4me3 (C) and total H3 (D) at the same genes. Cells were grown at 42°C (N), primed for 30 min at 57 °C (P), or primed followed by a 24 h lag phase (PL). NL denotes unprimed controls harvested in parallel. ChiP-pPCR values are shown as % input normalized to the control locus *CMB054C*. Mean ± SEM, n = 4 biological replicates. One-way ANOVA followed by a Tukey’s post-hoc test (α = 0.05); different letters indicate significance with p < 0.05. **E.** Relative expression of *CMJ099C-CMJ102C* across memory conditions in WT and E(z) determined by RT-qPCR. Two-Way ANOVA followed by Tukey’s HSD test, α = 0.05, n = 3, different letters indicate significance with p < 0.05.

**Figure EV5: HS memory in *E(z)*.**

**A.** Untransformed PSII quantum yields Fv/Fm previously primed (PLT) and non-primed (T) WT and *E(z).* Algae cultures were treated as described (Fig 1A) with a 24-h lag phase. Data show mean ± SD (n = 3 with three technical replicates per biological replicate). For transformed data Fv/Fm(PLT/T) see Fig 4B. **B.** Total numbers of up-(red) and downregulated (blue) genes identified in WT and *E(z)* at different time points of the thermotolerance experiment (Fig 1A, E). T and PLT samples were harvested after 15 min at 60 °C. Genes were considered significant DEGs if pl = -log(padj) x |L2FC| > 1.5 (n = 3 with three technical replicates per biological replicate).

---

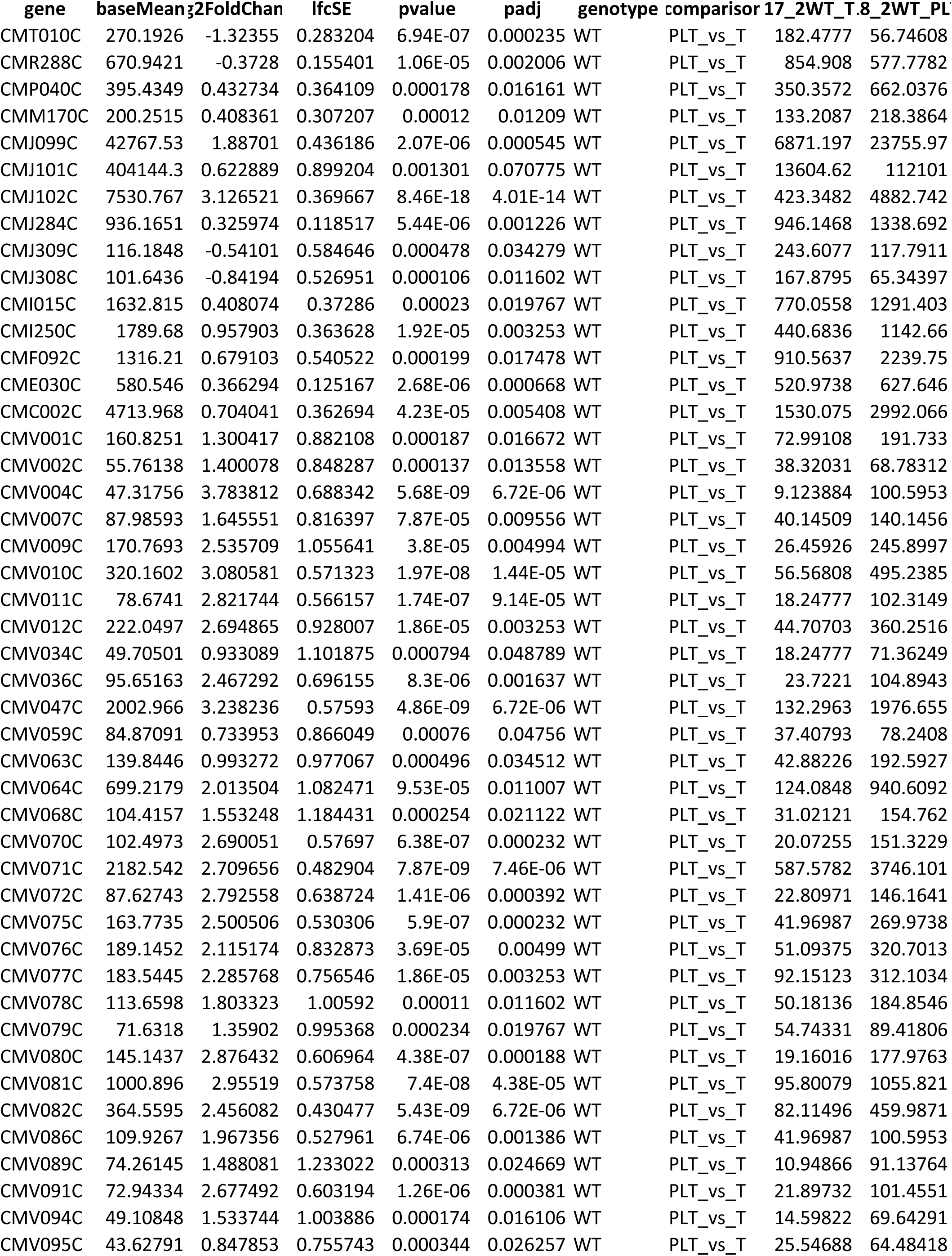
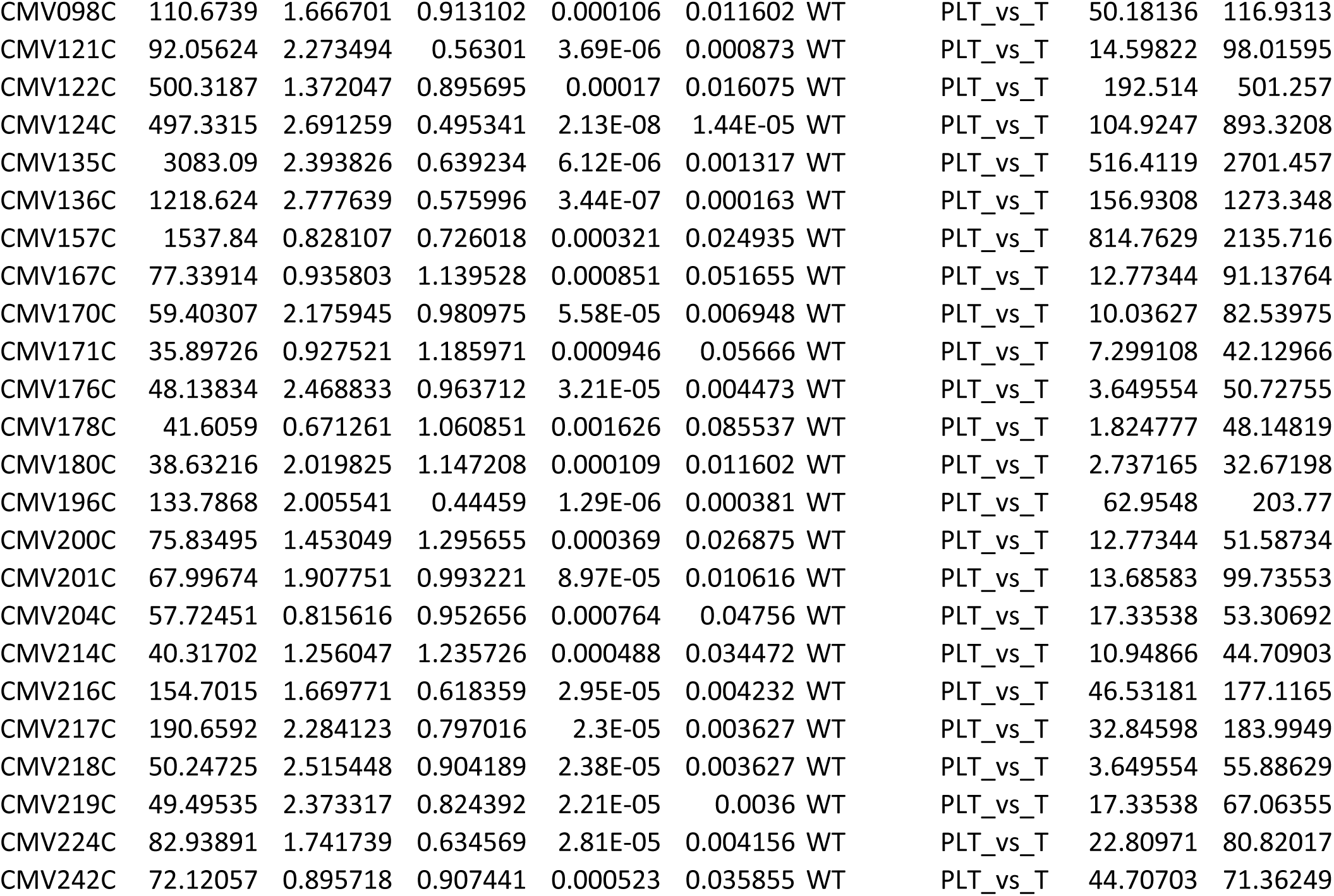

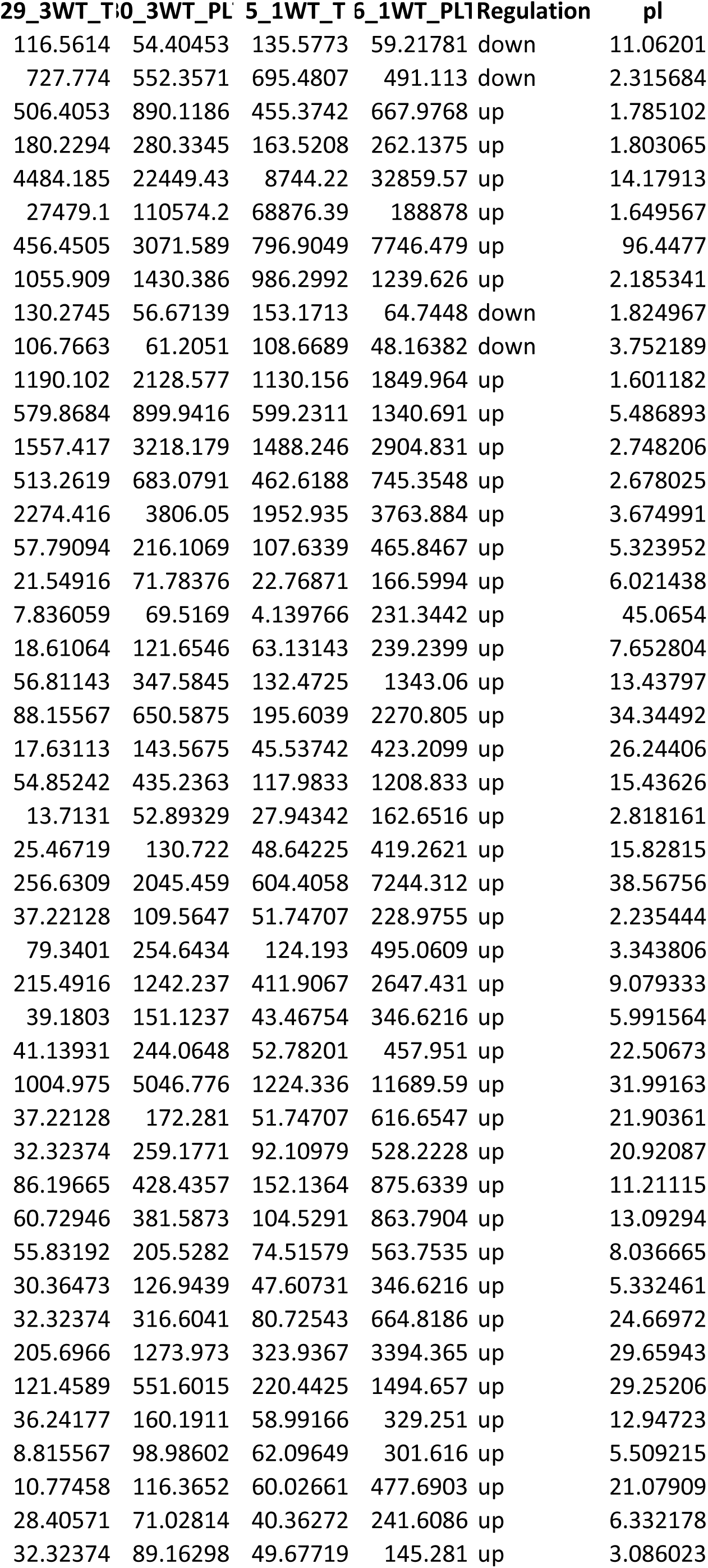
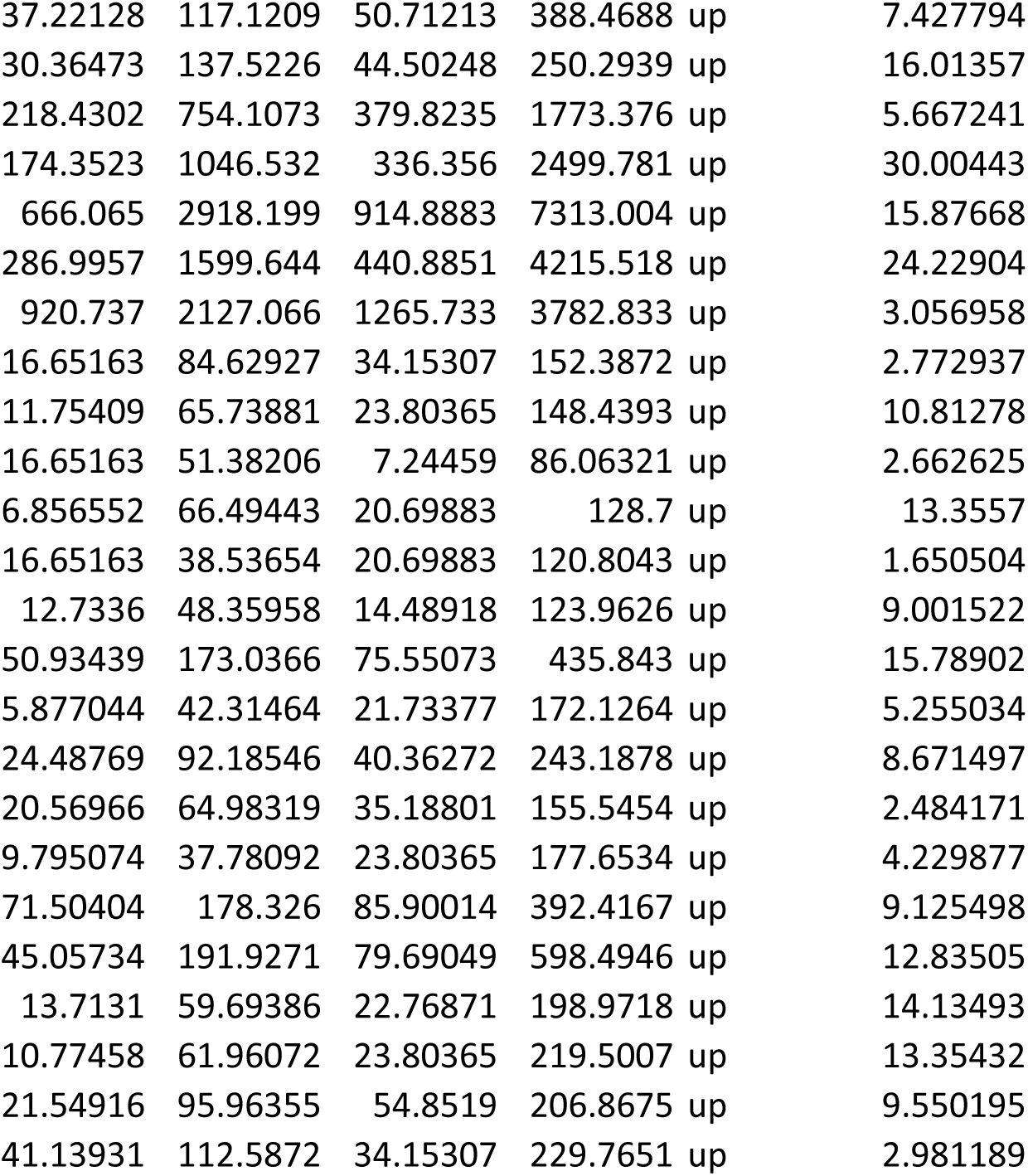

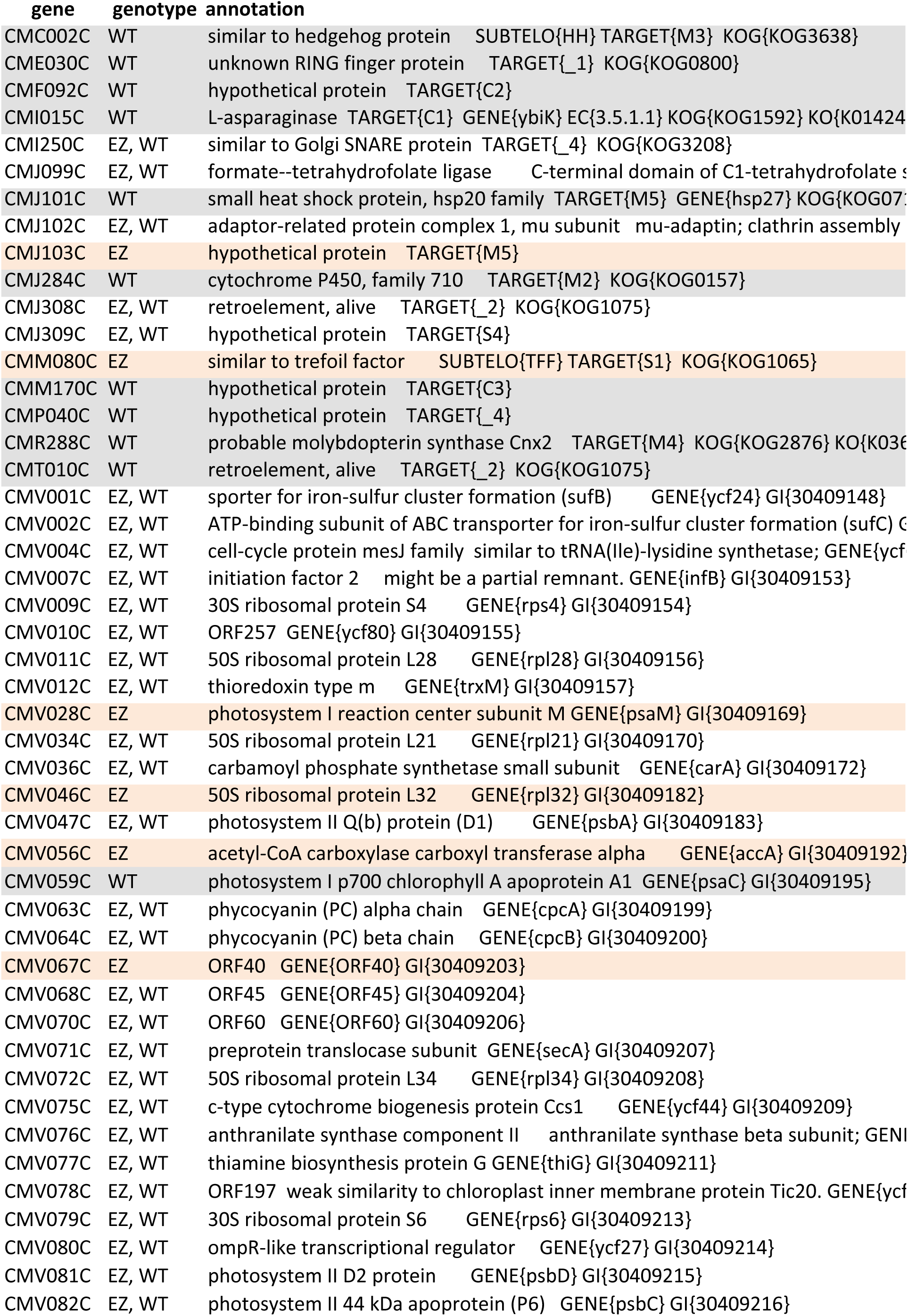
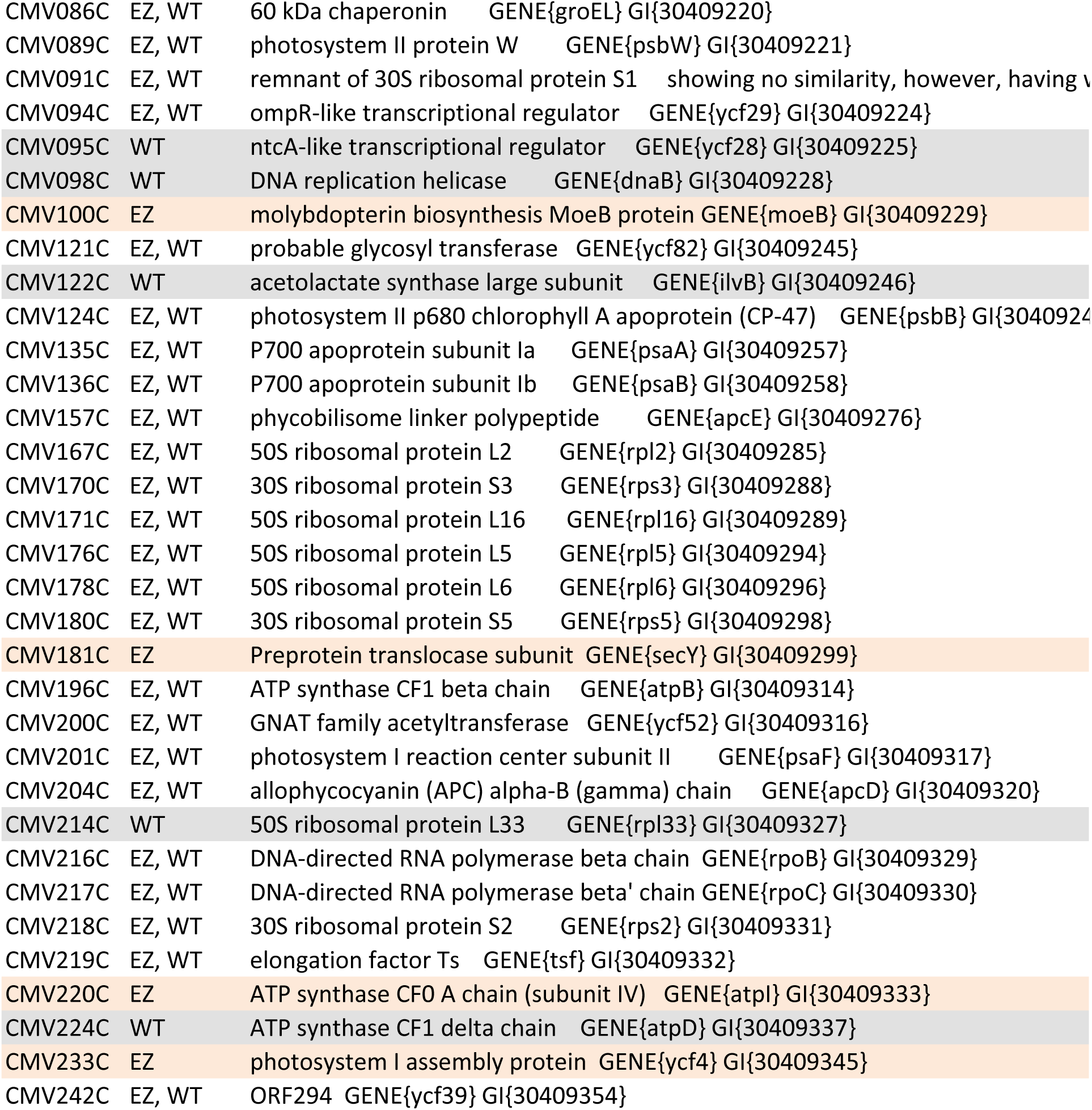

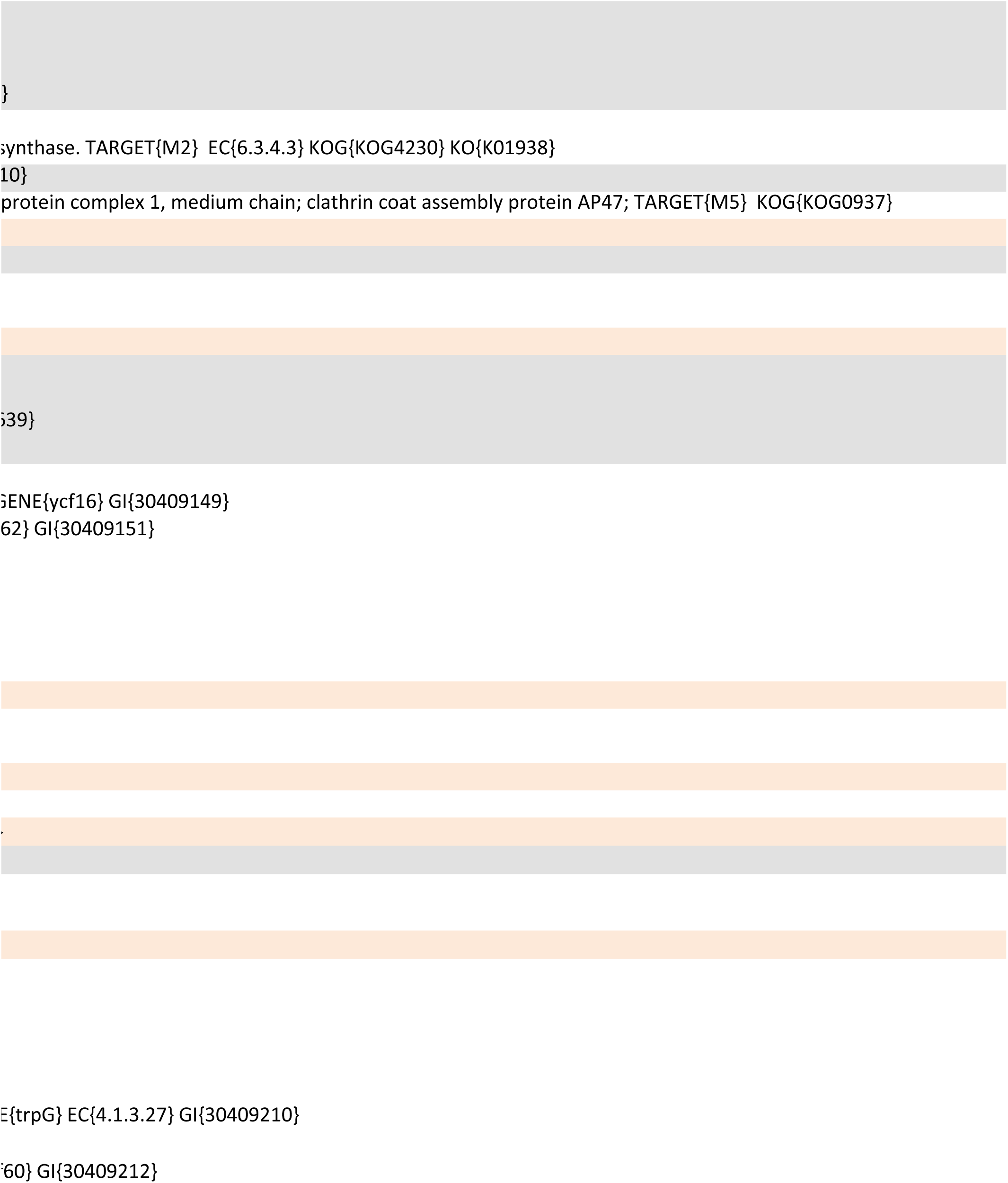
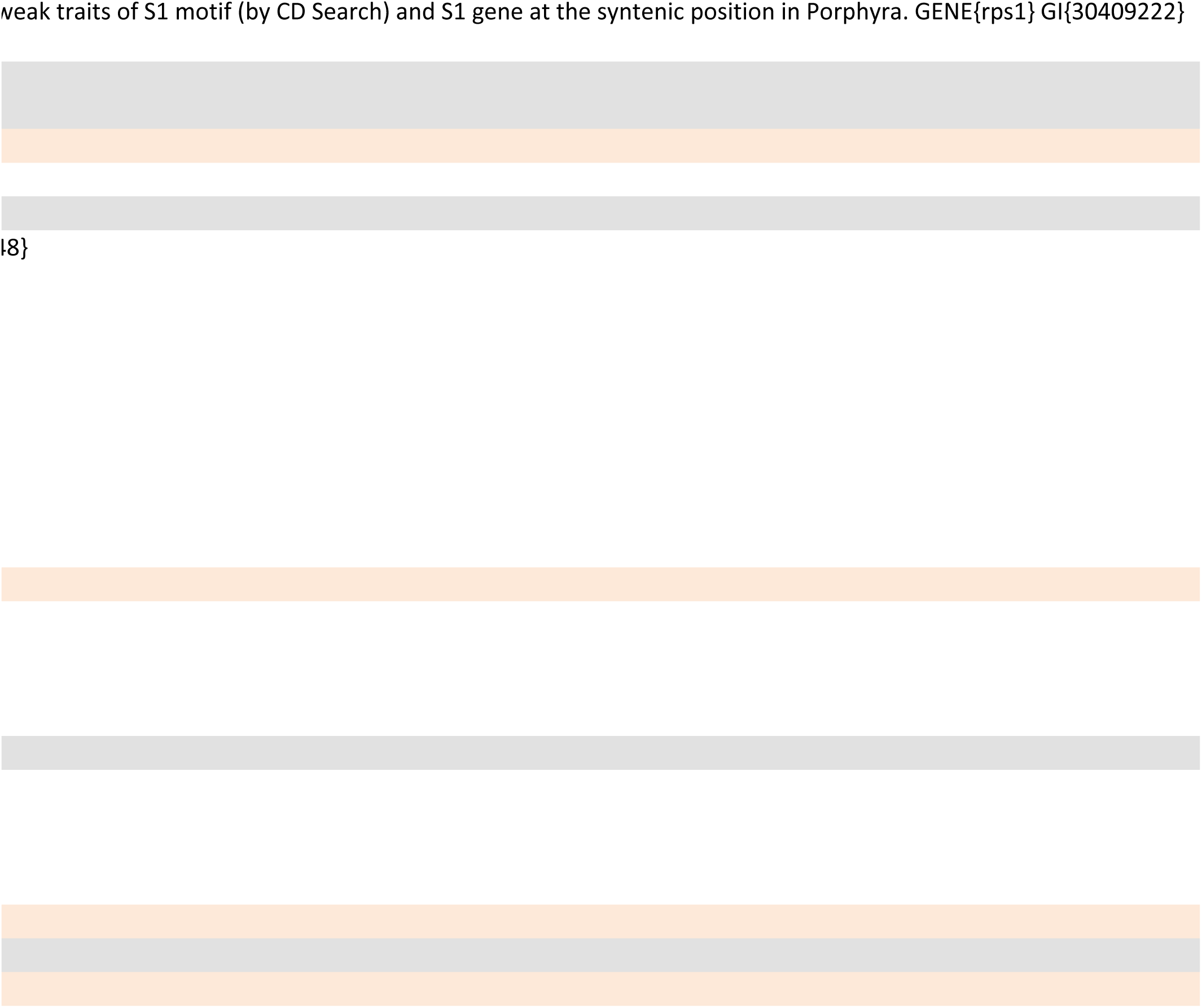

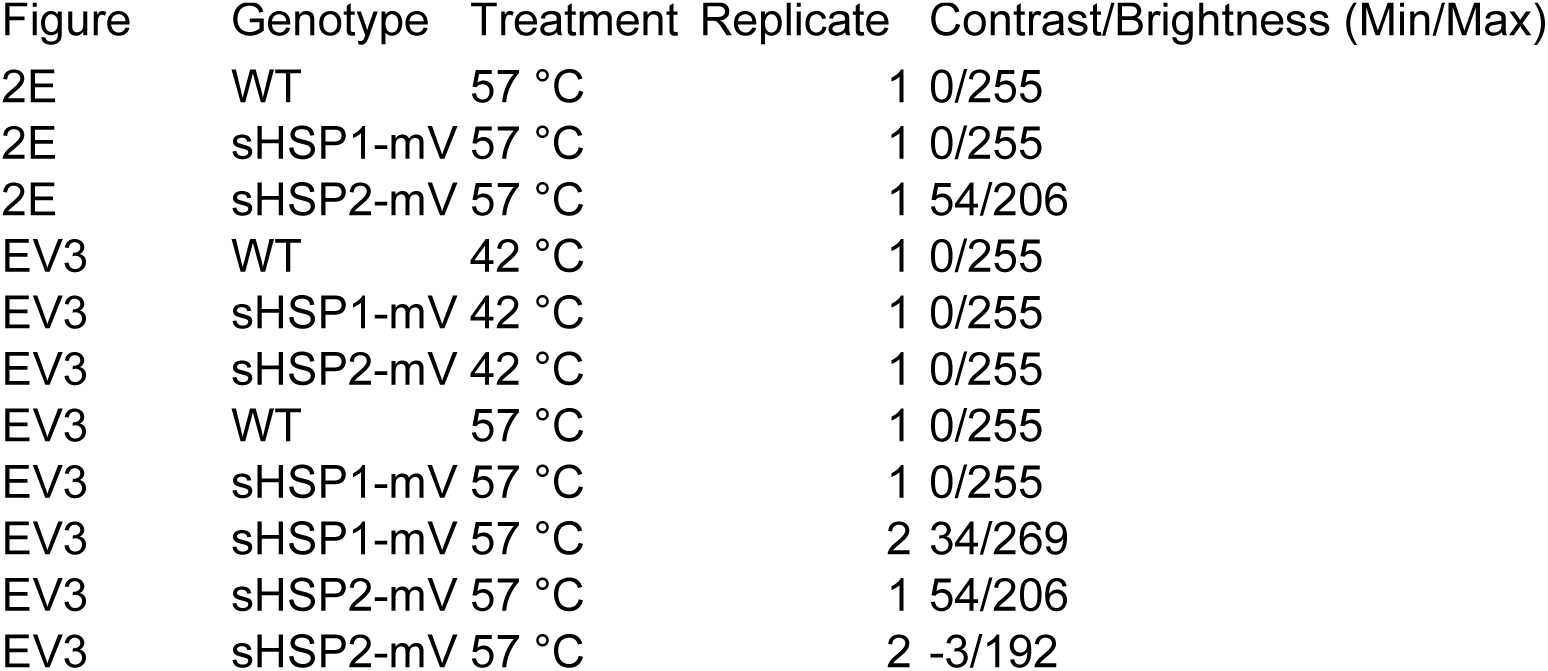

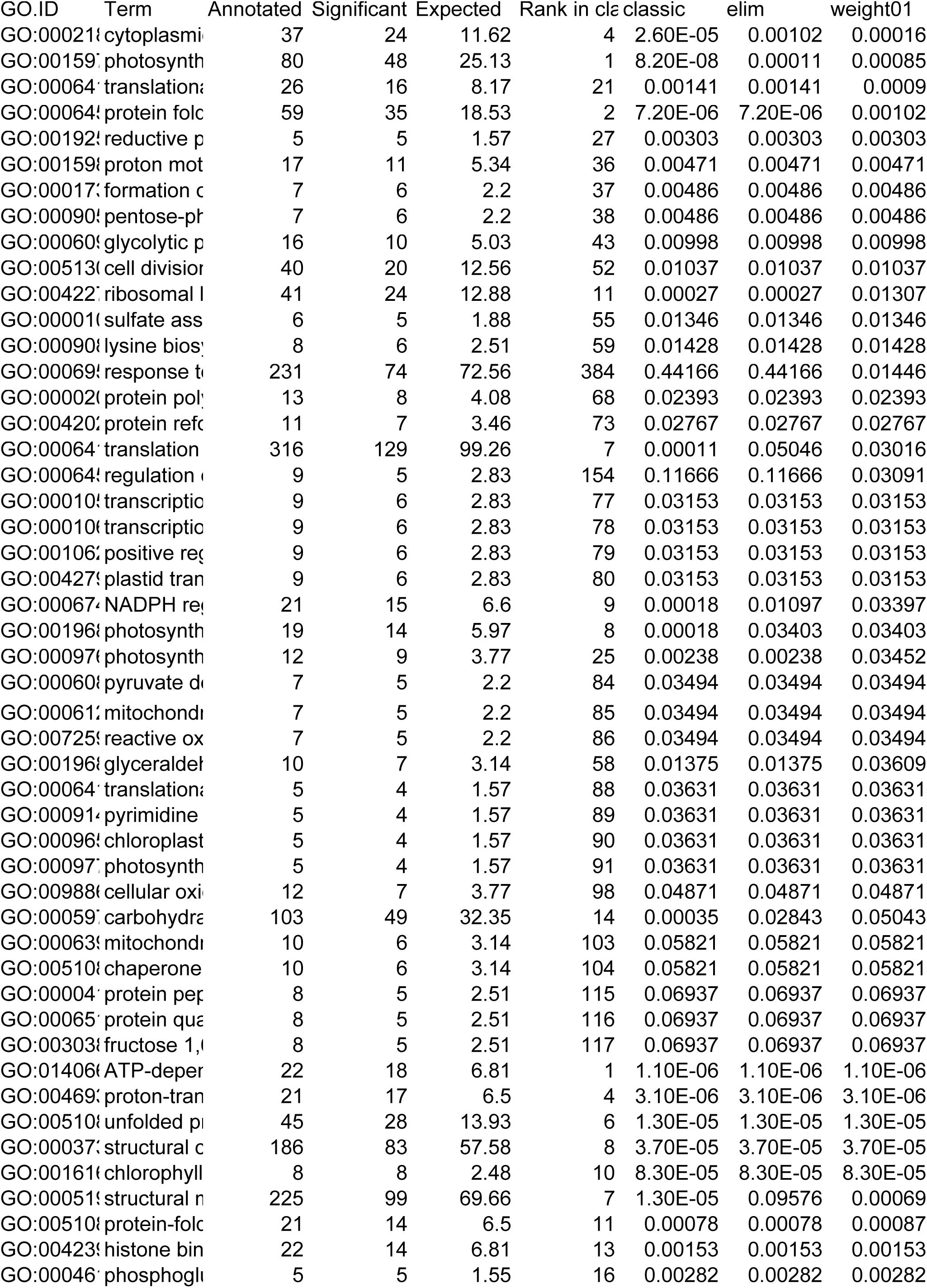
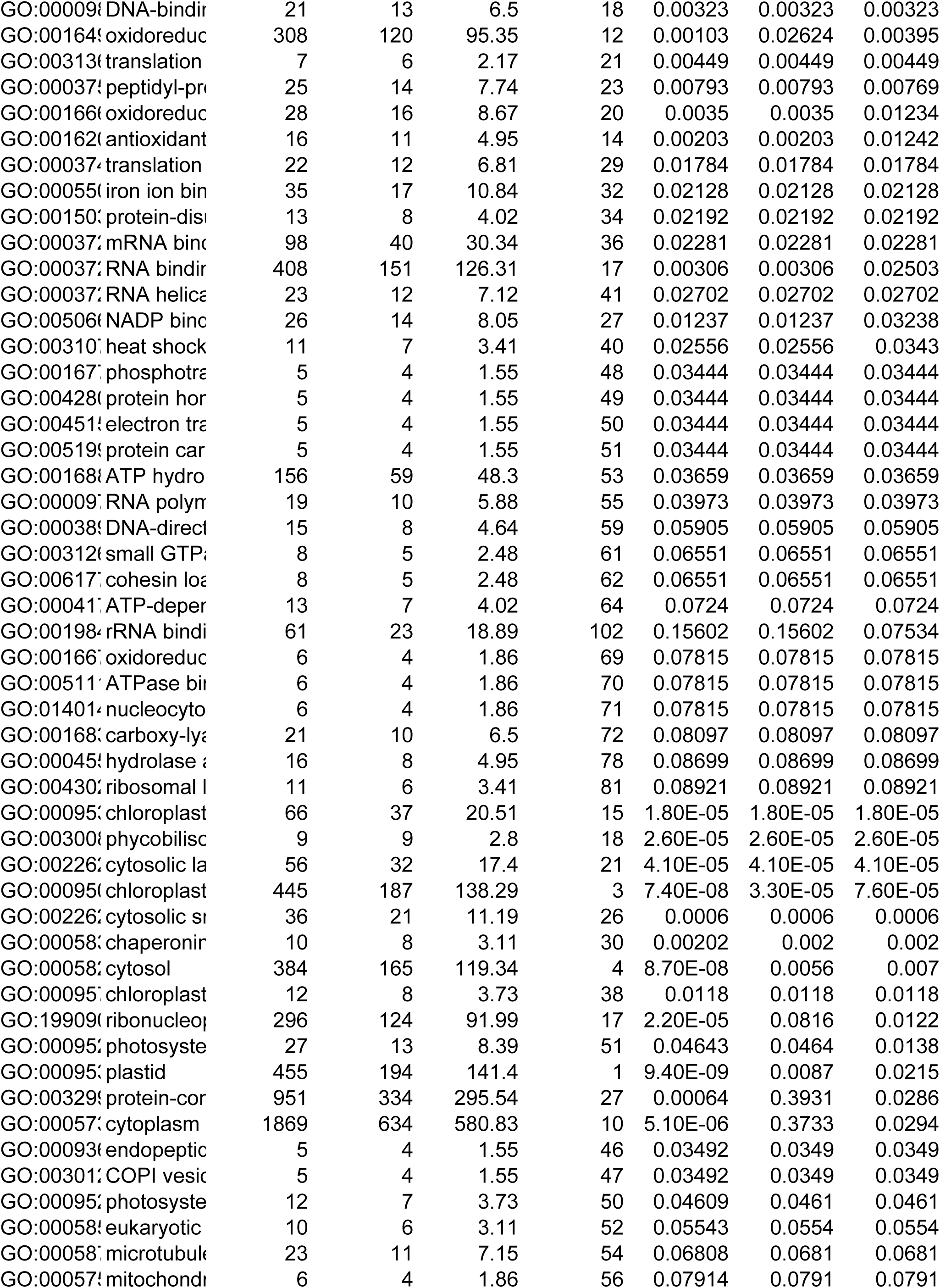
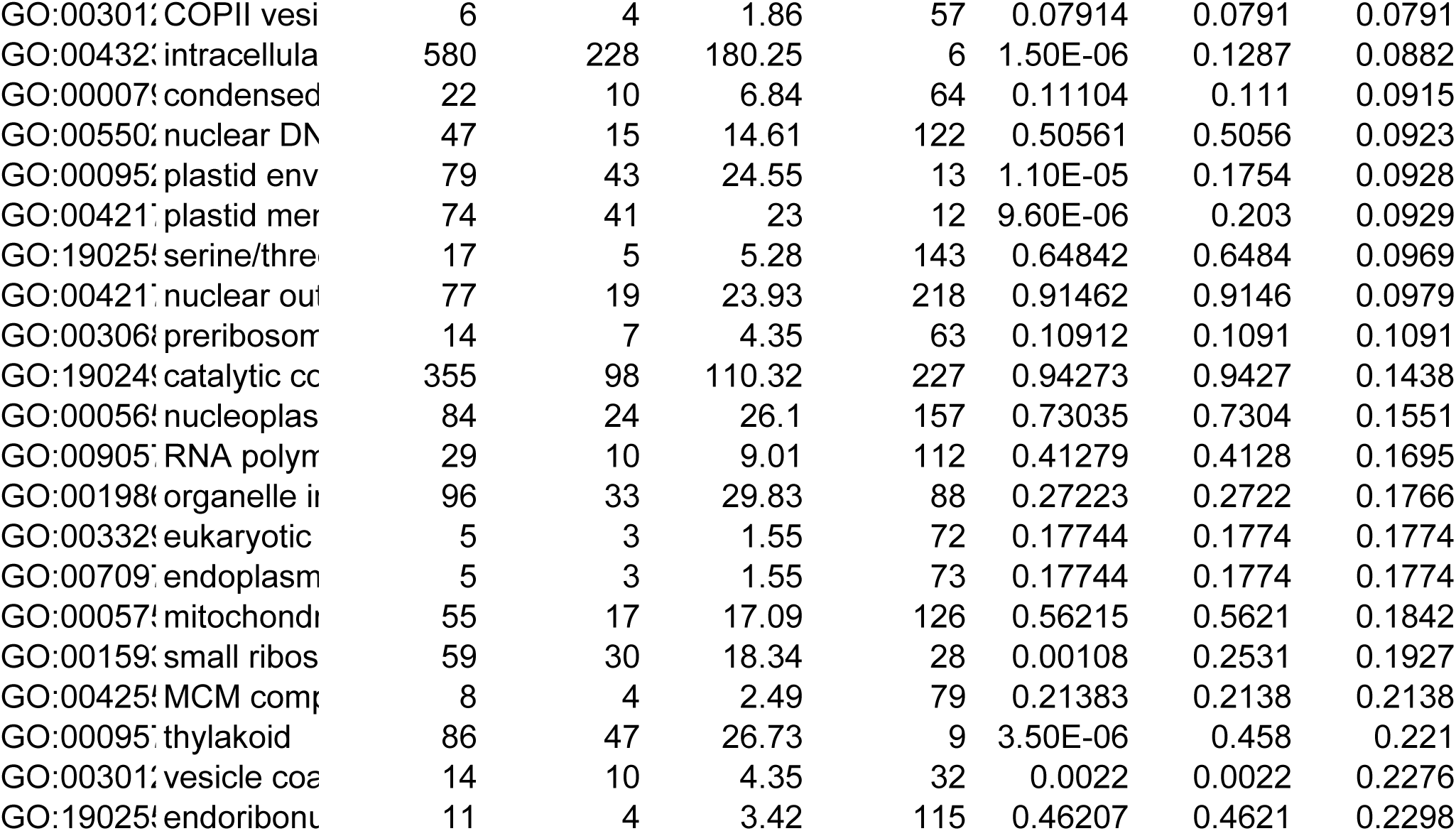

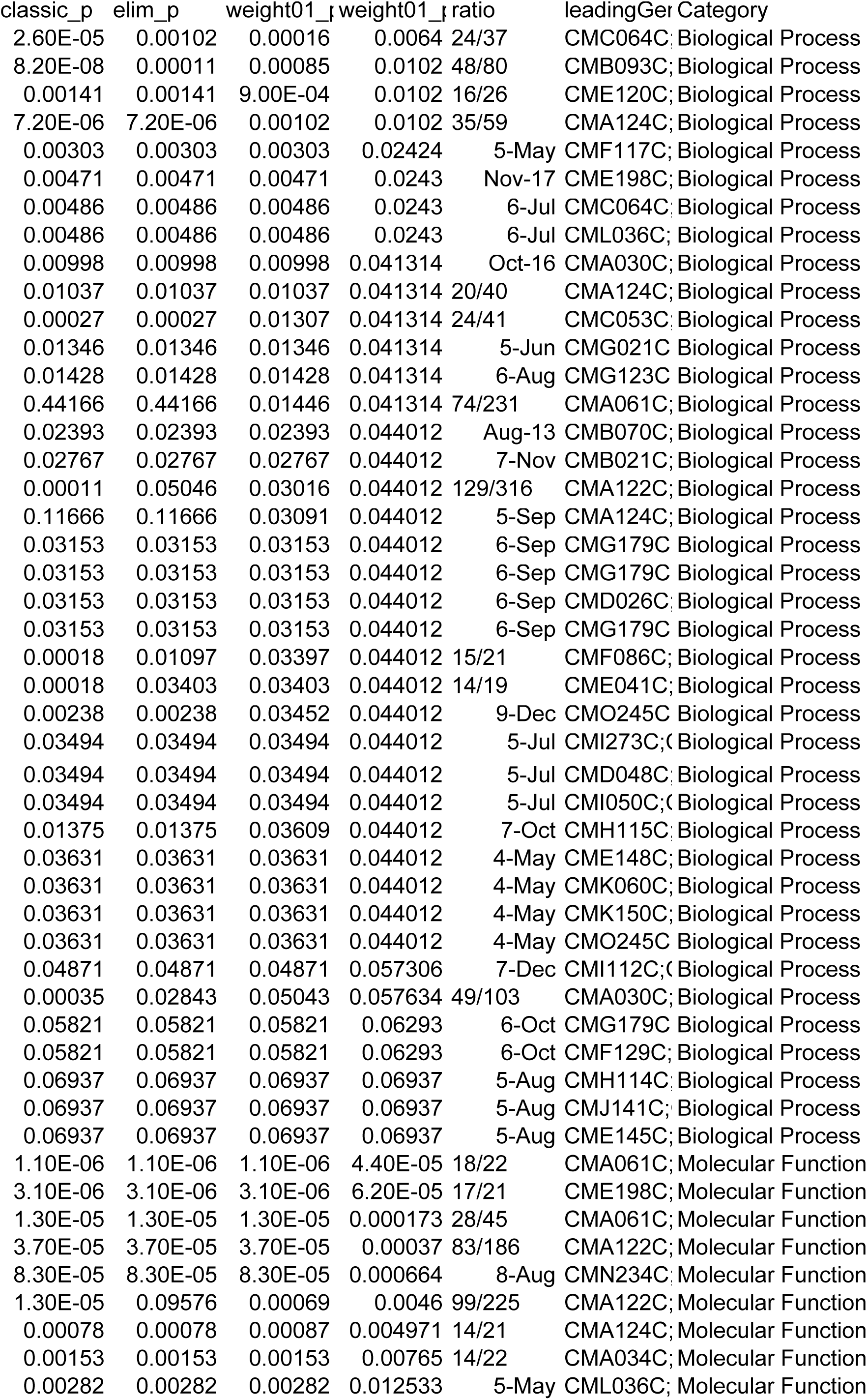
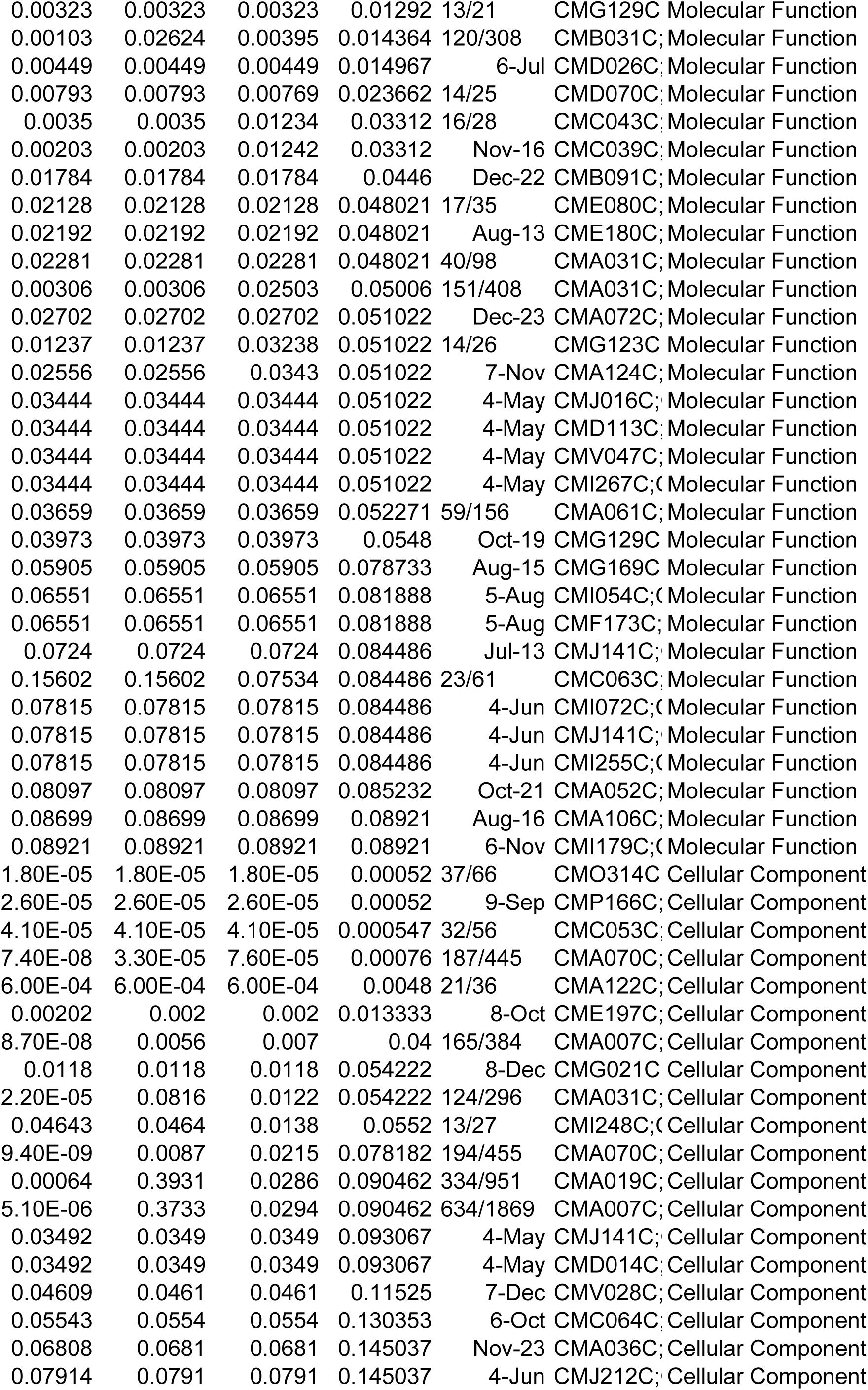
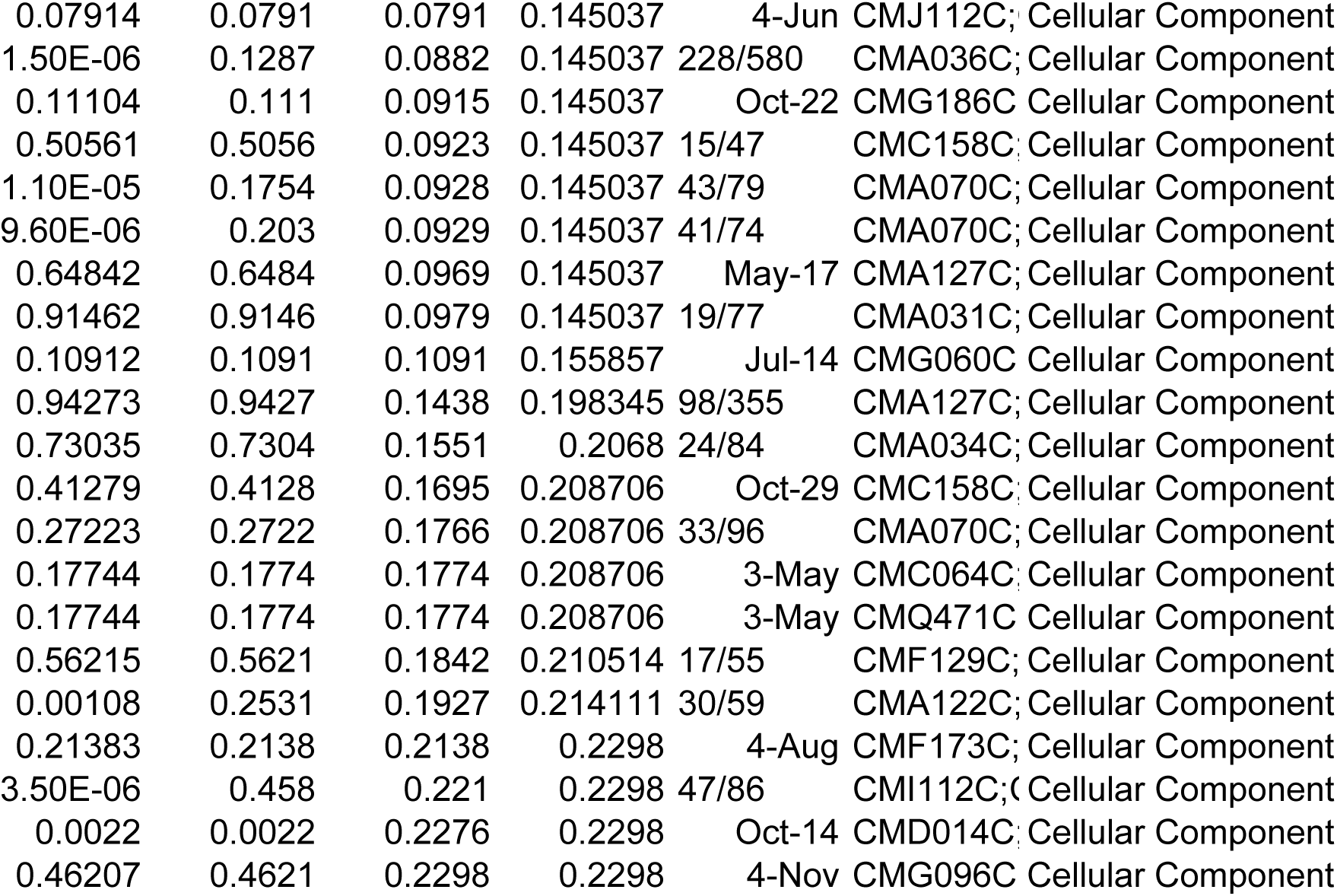

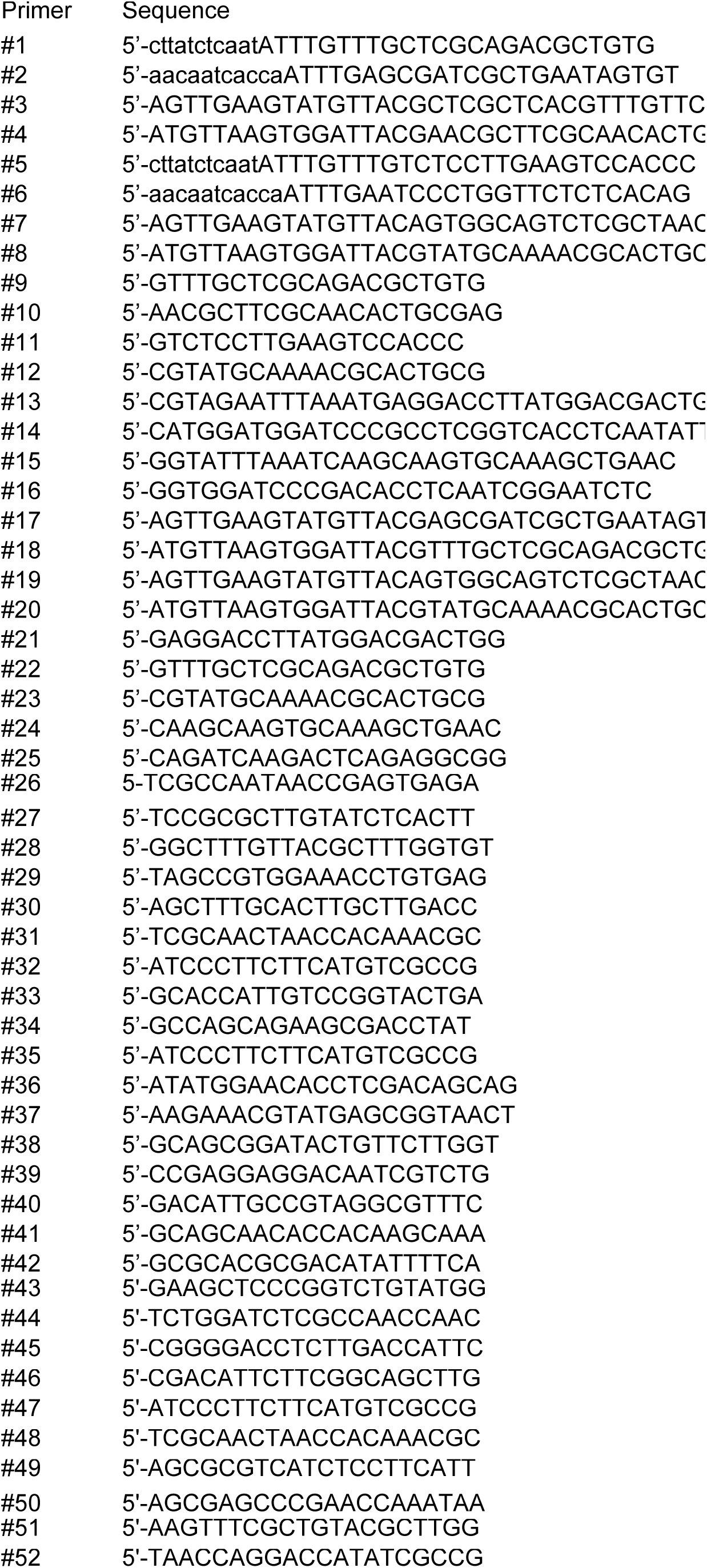
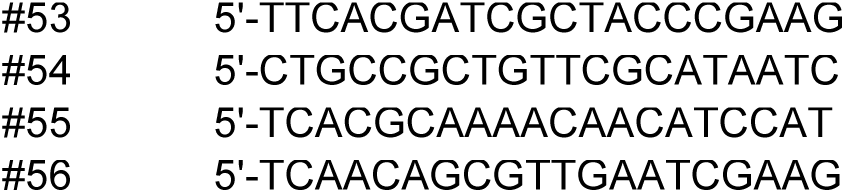

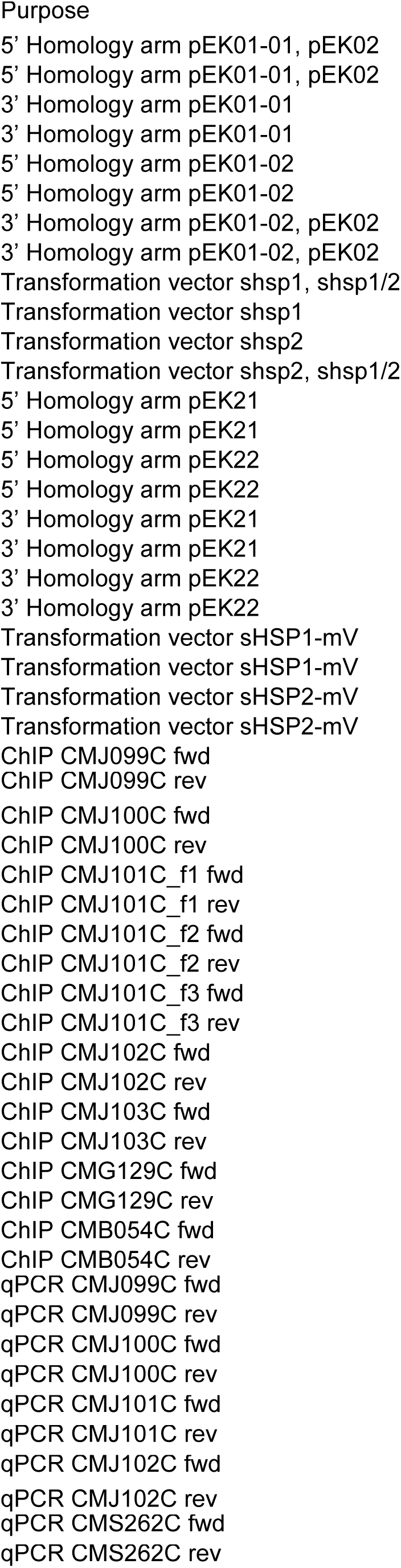
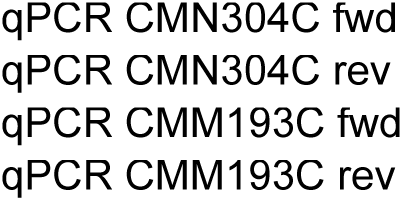

